# An increase in atypical petal numbers during a shift to autogamy in a coastal sand verbena and potential evolutionary mechanisms

**DOI:** 10.1101/2021.01.03.425117

**Authors:** Eric F. LoPresti, James G. Mickley, Addison Darby, Christopher G. Eckert, Michael Foisy, Cecilia Girvin, Sierra Jaeger, Katherine Toll, Alyson Van Natto, Marjorie G. Weber

## Abstract

**Premise:** In plants, meristic traits, such as petal and sepal numbers, are usually considered invariant within taxa, yet certain species consistently exhibit great variability in these traits. The factors contributing to “atypical” counts are not well-known, published hypotheses include relaxation of pollinator selection, inbreeding, and hybridization, among others. The sand verbenas, *Abronia* (Nyctaginaceae), usually have five perianth lobes (‘petals’), yet certain taxa exhibit marked departures from this norm.

**Methods:** Here we integrate an analysis of images from community science data (iNaturalist) and common garden experiments to evaluate a comprehensive set of adaptive and nonadaptive explanations for the production of these ‘atypical’ flowers across an evolutionary transition from xenogamy (outcrossing) to autogamy (selfing) in the coastal sand verbena *Abronia umbellata*.

**Key results:** The shift to autogamy in this lineage correlated with a higher frequency of atypical flowers from ~7% to ~20% and a significant reduction in mean petal number per inflorescence. Autogamous success did not change with petal number, and neither hybridization or up to three generations of inbreeding consistently increased production of atypical flowers or decreased mean petal number, all in contrast to previously-published hypotheses. In contrast, intra-inflorescence, inter-plant (intra-population), inter-population, and inter-variety comparisons demonstrated a correlation of reduced floral size with reduced petal number, suggesting correlated evolution due to a well-established relation between organ number and meristem size.

**Conclusions:** The reduction in petal number was probably a consequence of selection for smaller flowers associated with increased selfing. While we could not completely eliminate several alternative hypotheses, including a long-term history of inbreeding or relaxed selection on petal number constancy, those are less likely to explain the observed changes, though they may have contributed to the trend. In general, we develop a framework of hypotheses for evolutionary investigations of meristic variation in floral organs.

## INTRODUCTION

While the repeated modules that make up any plant vary substantially in phenotype (i.e. size, shape, number, chemical composition: Herrera 2009), vegetative traits are generally far more variable than floral traits. In plants, floral meristic characters such as petal number are easily quantified from fresh material, herbarium specimens, and even photographs. Because these characters are often considered fixed within species, genera, or families and, hence widely used for species identification and taxonomic classification, any departure from a ‘normal’ meristic character state within a taxon is of evolutionary, developmental, morphological, and taxonomic interest (Ellstrand, 1983; Ronse De Craene 2016). However, interspecific and intraspecific variation in the mean or variance of petal number is common in many families (e.g. Stark, 1918; Lowndes, 1931; Saunders, 1934; Roy, 1962; Huether, 1968; Schemske, 1978; Ellstrand, 1983; Lehmann, 1987; Shepard et al., 2005). Despite strong interest in meristic variation, critical examination of underlying evolutionary causes and adaptive significance is scarce (Ronse De Craene 2016).

Petal number variation can take two forms. Individual plants may differ from conspecifics in both their mean petal number as well as the variance in petal number. Given that most plants have a fixed base-petal number, the per-plant rate of production of flowers with atypical petal counts provides a metric for variation unaffected by whether petals are lost or gained. A suite of adaptive and non-adaptive hypotheses potentially explain shifts in either the mean or variance of petal number (Tables 1 & 2). The usual condition of remarkable invariance of petal number suggests a highly conserved developmental program (Endress 2001) perhaps as a result of pollinator selection for consistency in floral form mediated by innate preference (Stebbins, 1974; Herrera, 2009). While such pollinator selection has long been hypothesized (e.g. Leppik, 1953), supporting evidence remains elusive. Bees but not moths can ‘count’ petals (e.g., Leppik, 1953; Golding et al., 1999; Mickley, 2017). Mickley and Schlichting (2018) indirectly tested the pollinator consistency hypothesis by comparing outcrossing vs. autogamous species. If pollinators maintain a fixed petal number, nondirectional variation in petal number would be more frequent in autogamous than outcrossing species due to relaxed selection on petal number. In contrast to this expectation, atypical flowers were not more frequent in selfing vs. closely-related outcrossing species of Polemoniaceae.

**Table 1:** Hypotheses, and predictions thereof for both atypical petal counts and mean petal number tested in this paper.

| Hypotheses from literature tested here: | Predictions for *Abronia umbellata* | Specific prediction |
| --- | --- | --- |
| (H1) Loss of petals increases autogamous success | In *A. u.* var. *breviflora*, fewer petalled-flowers will have higher reproductive success in greenhouse | Negative correlation of petal number with seed set |
| (H2) Inbreeding increases developmental instability | (1) *A. u.* var. *breviflora* (autogamous) will have increased atypical petal morphs (2) Repeated inbreeding in *A. u. breviflora* will increase rates of atypical petal morphs | Increase in variance, agnostic to mean |
| (H3) Incompatibilities due to hybridization increase atypical petal morphs | (1) Artificially-created hybrids will have increased atypical petal morphs<br>(2) *A. u.* var. *umbellata* (xenogamous), which hybridizes with congeners commonly in the field, will have an increased atypical petal morphs in field compared to autogamous variety | Increase in variance, agnostic to mean |
| (H4) Correlated evolution with other traits (here: flower size) | *A. u.* var. *breviflora* (smaller flowered) will have fewer mean petals<br>Within each variety, individuals with smaller flowers will have fewer mean petals<br>Within each variety, populations with smaller flowers will have fewer mean petals | Decrease in mean, agnostic to variance |
| (H5) Environmentally-driven | A correlation between phenology and atypical petal morphs for each variety<br>A correlation between latitude and atypical petal morphs for each variety | Change in variance, agnostic to mean<br>Change in variance, agnostic to mean |
| (H6) Ontogenetically-driven | A correlation between age of plant and atypical petal production for each variety | Change in variance, agnostic to mean |

**Table 2:**
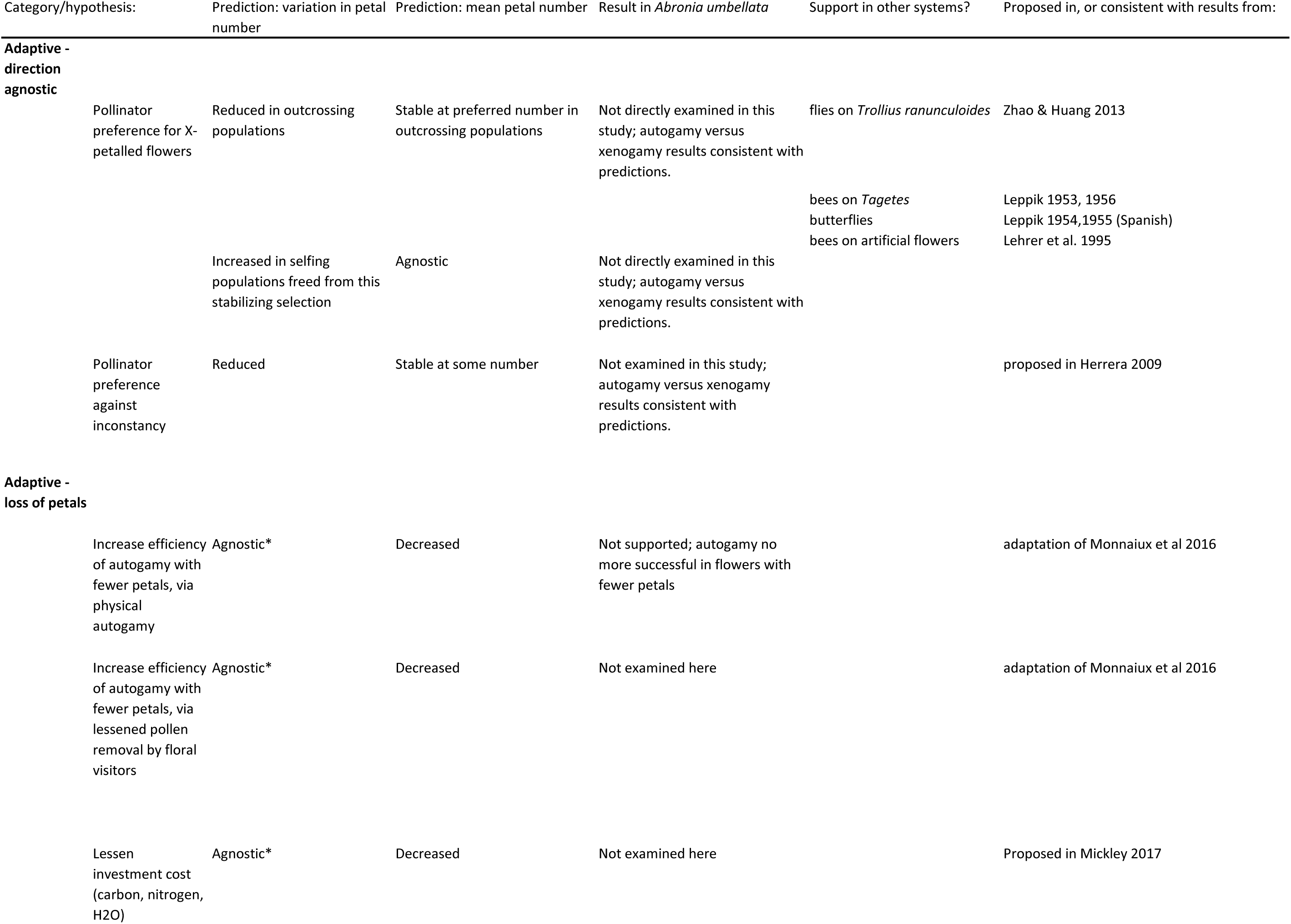
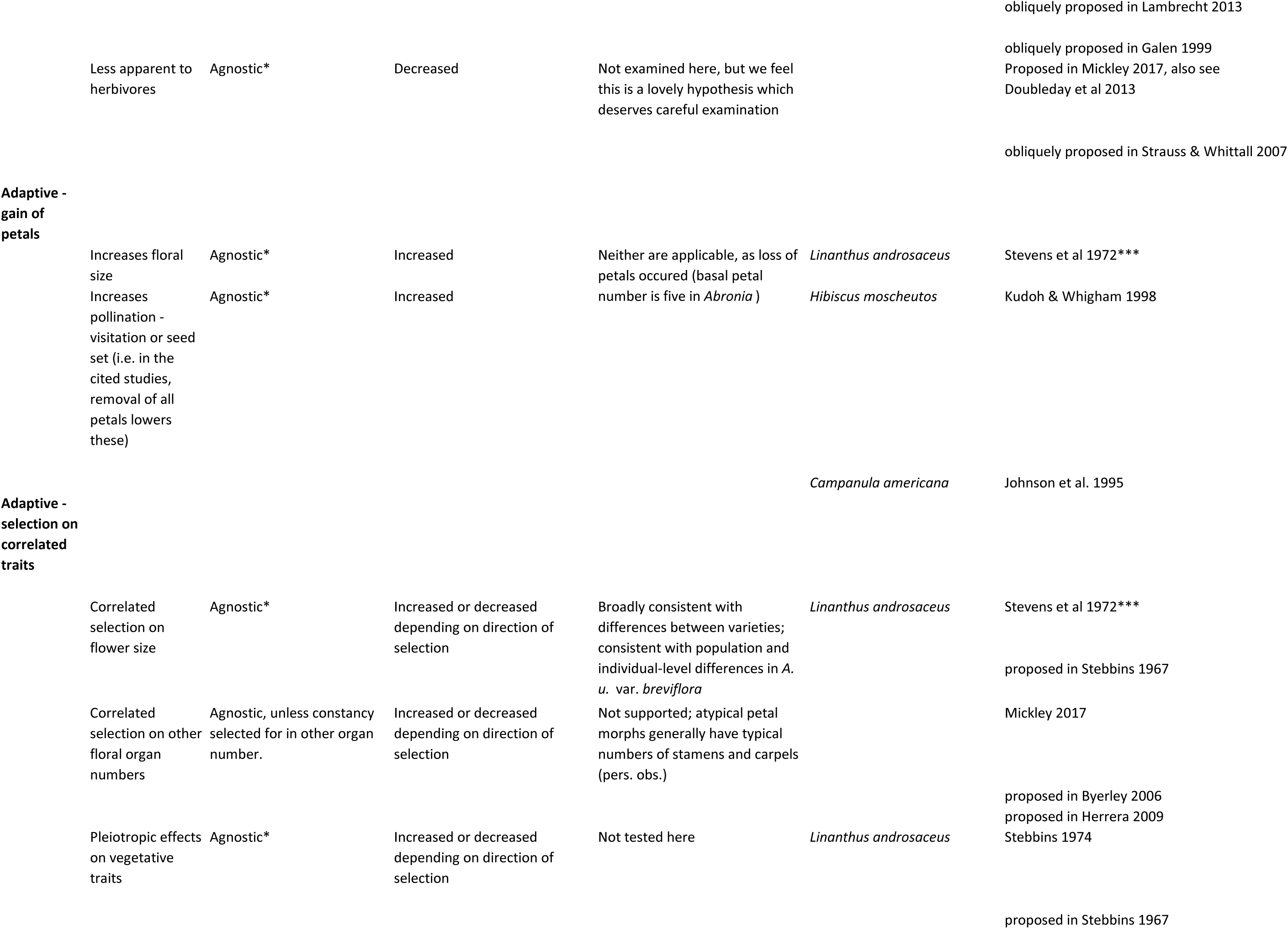
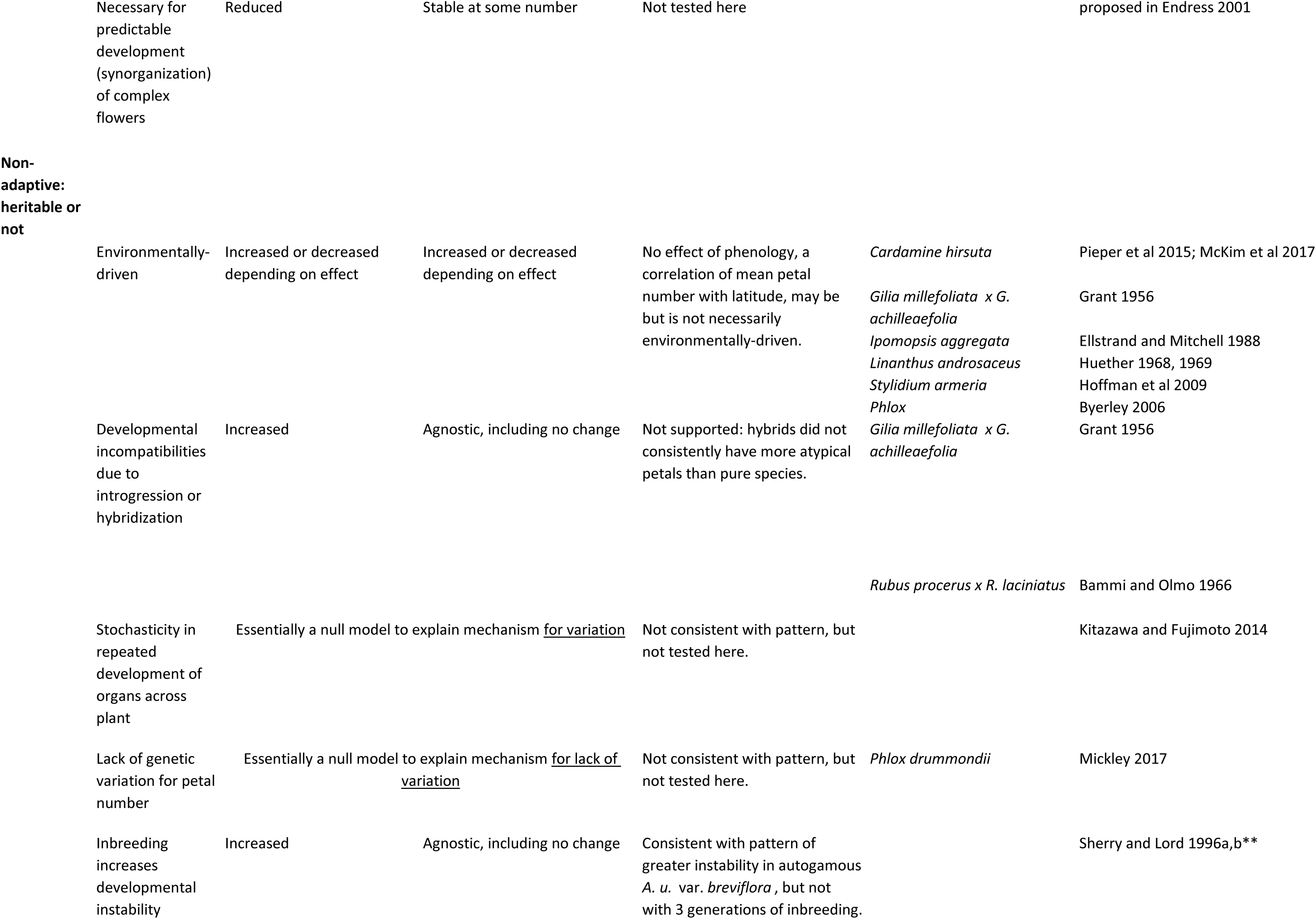
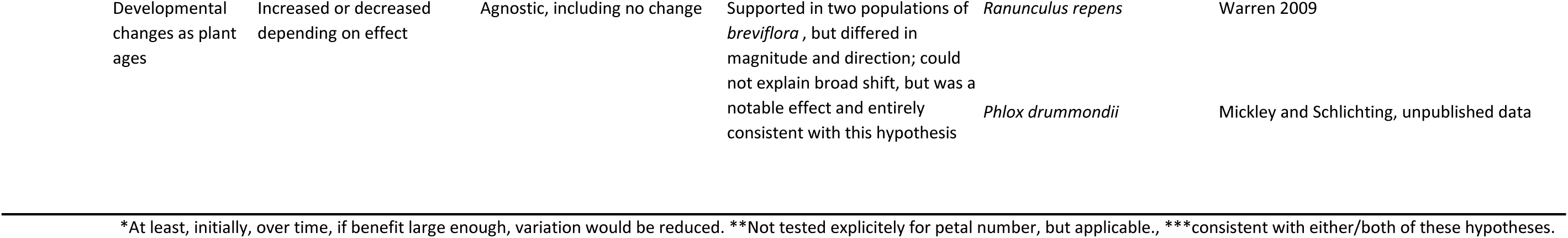
A broader set of evolutionary and ecological hypotheses for atypical petal morphs, with predictions for *Abronia umbellata*, and references.

The evolutionary transition to selfing may also be associated with a directional loss of petals, independent of relaxed pollinator selection if petals are costly (e.g., Galen, 1999; Strauss & Whittall, 2007; Lambrecht 2013) or if loss of petals facilitates self-pollination (Monnaiux et al 2016). For example, Monnaiux et al (2016) hypothesize that petal loss in *Cardamine hirsuta* slows bud opening and promotes more effective selfing (Table 1: H1). A later paper on the same system found that a genotype with fewer petals outcrossed less frequently in the field but did not differ in seed set in the lab (Monnaiux et al., 2018). Based on this work, we hypothesize two nonexclusive mechanisms potentially underlying their hypothesis. The first is that by delaying opening, reduced petal number would result in less pollen removed or deposited by pollinators, and therefore, outcrossing would be reduced. The second is that by delaying opening, the anthers and stigmatic surface would be in closer proximity for a longer time increasing the chance of pollen passively contacting the stigmatic surface. The former requires a field test with natural pollinators; the latter can be tested in the laboratory (Table 1: H1).

Non-adaptive evolutionary or phenomenological hypotheses also explain increased petal number variation. For example, inbreeding itself, independent of relaxed pollinator-mediated selection could reduce the developmental stability of flowers thereby increasing production of flowers with atypical petal numbers (agnostic to the petal loss or gain) (Table 1: H2). Hybridization may also disrupt developmental canalization due to genetic incompatibilities (e.g. outbreeding depression, Dobzhansky-Muller incompatibilities, Bachman et al., 1981; Vlot et al 1992; Pelabon et al., 2003), leading to higher production of atypical floral morphs (Table 1: H3). Hybridization increased the frequency of flowers with atypical petal numbers in certain intrageneric hybrids (Grant, 1956; Bammi and Olmo, 1966) and in others decreased this same metric (Choi et al. 2001; Bletsos et al. 1998). For example, hybridization between *Microseris* species with 5 vs. 10 pappus parts dramatically increased pappus number variation in the F1-F3 generations (Bachmann et al., 1981). We have no a priori reason to expect a directional change in mean, instead an increase in variance. Notably, we do not expect a uniform directional change in mean if incompatibilities due to hybridization destabilizes petal number, we likely would observe an increase in variance. Alternately, hybridization could reduce the variance in petal number, as found by Choi et al. (2001).

All the hypotheses discussed so far assume a genetic component to variation in the mean and/or variance of petal number. Heritability of genetic control of mean or variance in petal number has been demonstrated in many species (Stubbe, 1952 in Grant, 1956; Grant, 1956; Alpi et al., 1968; Katsuyoshi and Harding, 1969; Stevens et al., 1972; Agren and Schemske, 1995; Delesalle and Mazer, 1995; Mazer et al., 1999; Byerley, 2006, Pieper et al., 2015; Roman et al., 2015; Monnaiux et al., 2016, 2018; Mickley 2017). This heritability suggests that selection could act directly on petal number, but also could act on genetically-linked traits (e.g. Table 1:H4). Strong evidence links organ number, including petal number, with the size of floral meristem preceding organ initiation (Stevens et al 1972; Green 1992; Running et al 1998, Fletcher 2001, Barrero et al 2006; Ronse De Craene 2016, also proposed in Stebbins 1967, Levy 1997). Experimentally treating plants with growth inhibitors to reduce meristem size (and subsequent floral size) often reduces floral organ number, the opposite treatment often increases it (reviewed in Green 1992). Artificial selection for increased and decreased organ numbers alters meristem size in the same direction (Stevens et al 1972). Several mutations at a single loci that alter meristic size produce correlated changes in floral organ number in *Arabidopsis* and *Solanum* (Running et al 1998, Fletcher 2001, Barrero et al 2006). Therefore, natural selection favoring smaller flowers for whatever reason (e.g. selfing, or reducing damage from flower-visiting herbivores, Doubleday et al 2013), may indirectly reduce petal number, leading to individuals or populations with smaller flowers having fewer mean petals per flower.

Finally, while petal number is often heritable, there is evidence for plasticity in petal number (Table 1: H5). Mean or variance in petal number may be affected by day length, temperature, herbivory, and with variation in the abiotic or biotic environment along latitudinal and elevational gradients (Stubbe 1952 in Grant, 1956; Grant, 1956; Roy, 1962; Huether 1968, 1969; Mathiason, 1982; Delesalle and Mazer, 1995; Tooke and Battey, 2000; Hoffman 2009; McKim et al 2017). Ontogenetic changes could also explain variation in certain cases (H6). Older clones of a buttercup (*Ranunculus repens*) had higher likelihood of producing extra petals, a result which Warren (2009) ascribes to the accumulation of somatic mutations over time, but this result could also reflect other age-related factors, such as hormonal changes.

The various hypotheses proposed for petal number variation have not been addressed in a single system using broad sampling across species’ ranges. Petal number can be scored on live plants, herbarium specimens, or photographs from rapidly growing georeferenced photographic databases from community science initiatives (e.g. iNaturalist, CalFlora, etc). Therefore, these databases represent an exciting complement to herbarium and other natural history collections. Here, we use approaches spanning spatial scales and involving both field and common garden experiments to address several proposed hypotheses for atypical petal numbers (Table 1) using the coastal sand verbena *Abronia umbellata*. In particular, we investigate phenotypic variation across a natural transition in mating system where southern populations (var. *umbellata*) are outcrossing, the ancestral condition in this clade), and northern populations (var. *breviflora*) are selfing.

## MATERIALS AND METHODS

### The geographic range of Abronia umbellata and its varieties

*Abronia umbellata* grows on beaches and coastal dunes – never more than a few hundred meters inland - from British Columbia, Canada, south to Baja California, Mexico. It has two, geographically separated, genetically-distinct varieties across its range (Greer, 2016; Van Natto, 2020). The southern variety (*A. u. umbellata*) is pollen-limited and self-incompatible, and therefore requires pollinator visitation, primarily by moths (Doubleday et al 2013; Doubleday and Eckert 2018). In contrast, the northern variety (*A. u. breviflora*) is self-compatible, highly homozygous for genome-wide SNPs (Van Natto and Eckert 2022) and probably nearly completely self-fertilizing in nature (Doubleday et al., 2013; Doubleday and Eckert, 2018; LoPresti, pers. obs.). Transition from outcrossing to selfing has occurred once and recently in this species (Greer, 2016). Almost all *Abronia* species are self-incompatible (Tillett, 1967; Galloway, 1975; Williamson et al., 1994; Darling et al., 2008), including those closely related to *A. umbellata* (LoPresti, unpublished data), further, there are very few unique SNPs in populations of var. *breviflora*, and thus, any shift in production of atypical flowers in *A. u.* var. *breviflora* occurred with a transition to autogamy (Doubleday et al., 2012; Greer et al. 2022; Van Natto and Eckert 2022). Hybridization occurs commonly in *Abronia*, especially in the xenogamous *A. u.* var. *umbellata,* which hybridizes naturally with *A. latifolia, A. maritima, A. villosa, A. gracilis*, and probably *A. pogonantha* (Tillett, 1967; Johnson, 1977; Van Natto, 2020; LoPresti, pers. obs.). While Tillett (1967) hypothesized that *A. u.* var. *breviflora* may be the result of a hybridization event between *A. umbellata* var. *umbellata* and *A. latifolia*, using genetic methods, Van Natto (2020) found that hypothesis extremely unlikely. Because of locally-common hybridization in nature (Pimental 1981), through limited introgression (Van Natto and Eckert 2022), hybridization could alter petal counts in certain regions.

### Broad survey of petal number across the range of Abronia umbellata (H2/H3/H4/H5)

To quantify petal number variation in the field, we counted petals on flowers from iNaturalist photographic records from locations across the entire species’ range. (n=1096 “research grade” records as of 13-Jan-2020). Of these, 865 records had sufficient detail to be scored, comprising 1,808 inflorescences, and 11,428 individual flowers. For each inflorescence in the photograph with enough visibility to score, we recorded petal number of each flower, as well as record number, latitude, and the date of the record. For records where the location was obscured by iNaturalist, the latitude listed was taken, as the obscuring occurs within a 10 km radius, a relatively small distance compared to the range-wide scale of this study and therefore, this uncertainty should not affect our results. Because interdigitation between prostrate stems of closely-packed plants is common, we were unable to conclusively assign inflorescences to individual plants therefore, multiple inflorescences in the same photograph were scored separately, but with the iNaturalist record number the same; the record number was used as a random effect in analyses when necessary (as a rough proxy for individual). 18% of records were not scorable, as petals were not countable in the photos. An additional 3% of ‘research-grade’ records were also discarded because of misidentifications: one *Abronia maritima* was misidentified as *A. umbellata* and 28 records were obvious hybrids between *A. umbellata* and co-occurring congeners *A. maritima* or *A. latifolia*.

To determine whether mating system predicted atypical petal counts (i.e. not 5-lobed), we used a binomial mixed model (R 3.6.2: package *lme4* version 1.1-27.1, https://github.com/lme4/lme4/) of with typical, atypical petal number as a binary response variable, variety as a fixed effect and iNaturalist record number as a random factor. We did the same for mean petal number using a gaussian error-distribution. These models were each compared to null models without variety using likelihood ratio tests.

### Common garden petal counts (H2, H3, H4, H5)

To quantify variation in petal counts between outcrossing and selfing varieties while minimizing potential environmental variation, we did two greenhouse common garden studies. The first, at Michigan State University, involved 10 selfing (var. *breviflora*) individuals and six outcrossing (var. *umbellata*) individuals from seed collected by EFL or procured from the USDA germplasm resource information network (Supplementary Table 1). Plants were maintained in the greenhouse and laboratory in 8-18” pots, in a mixture of sand and potting soil (~1:1), under 12h of artificial light per day in addition to ambient light. Plants were watered every 3-5 days and fertilized (Miracle-Gro, 10:10:10) once a month. Plants were moved between the greenhouse and laboratory often, for all measurements. We counted petal number of flowers in mature inflorescences of every plant that flowered between July 2019 and May 2020.

### Effect of inbreeding and ontogeny on atypical floral production (H2/H6)

The second common garden experiment took place in a greenhouse at Oklahoma State University and involved a total of 99 selfing (var. breviflora) plants, from three populations and with three levels of inbreeding (wild-collected seed, first generation lab-selfed seed, second generation lab-selfed seed). The conditions were the same as the previous experiment (except for 16h light/day). For plants, we counted petals on all flowers in the first 20 inflorescences per plant, as well as several inflorescences per plant at the conclusion of all plants making their 20^th^ inflorescence (for those plants still flowering at that point); in total we scored 28244 flowers from 2182 inflorescences. Floral measurements (tube length, limb to top of anthocarp, floral face diameter, and longest axis across lobes) on a haphazardly chosen single edge flower per inflorescence were taken when time permitted (n=1349, 62% of scored inflorescences).

### Hybridization effects (H3)

To examine the petal numbers of hybrids, we scored all hybrids of *A. umbellata* available on iNaturalist (n=86), UC-Berkeley’s CalPhotos (n=4), in E.F.L.’s and A.V.N. personal photographs of plants in the field (n=6), and laboratory crossed plants in the common garden (n=37; Supplementary Table 1). Hybrids were identified by using morphological traits of floral color, inflorescence shape, leaf shape and leaf succulence (Tillett 1967; LoPresti & Van Natto, pers. obs.). For the iNaturalist data, using a binomial mixed model, we compared atypical petal production in pure species (from the broad iNaturalist data) to that of the hybrids, with record as a random effect, for each variety. For var. *umbellata*, which hybridizes with both *A. maritima* and *A. latifolia*, we used the hybrid cross as an additional fixed effect.

### Laboratory test of autogamy success with different petal numbers (H1)

To test the adaptive hypothesis that decreased petal number increases autogamous success, we measured covariance between seed set via autogamy and petal number of 597 individual flowers (597 flowers (417 5-petalled, 169 4-petalled, 6 3-petalled) during 7-November-2019 and 09-March-2020 in 58 inflorescences produced by selfing plants in the MSU common garden (Fig. 1: E, F). The six flowers with three petals and five which did not open were excluded from analyses. Because *Abronia* ovaries contain a single ovule we fit seed set as a binomial (seed/no seed) response with petal number as a fixed effect and plant and inflorescence (within plant) as random effects. To evaluate the effect of petal number, we compared this model to a null model with only the random effects using a likelihood ratio test.

**Figure 1.**
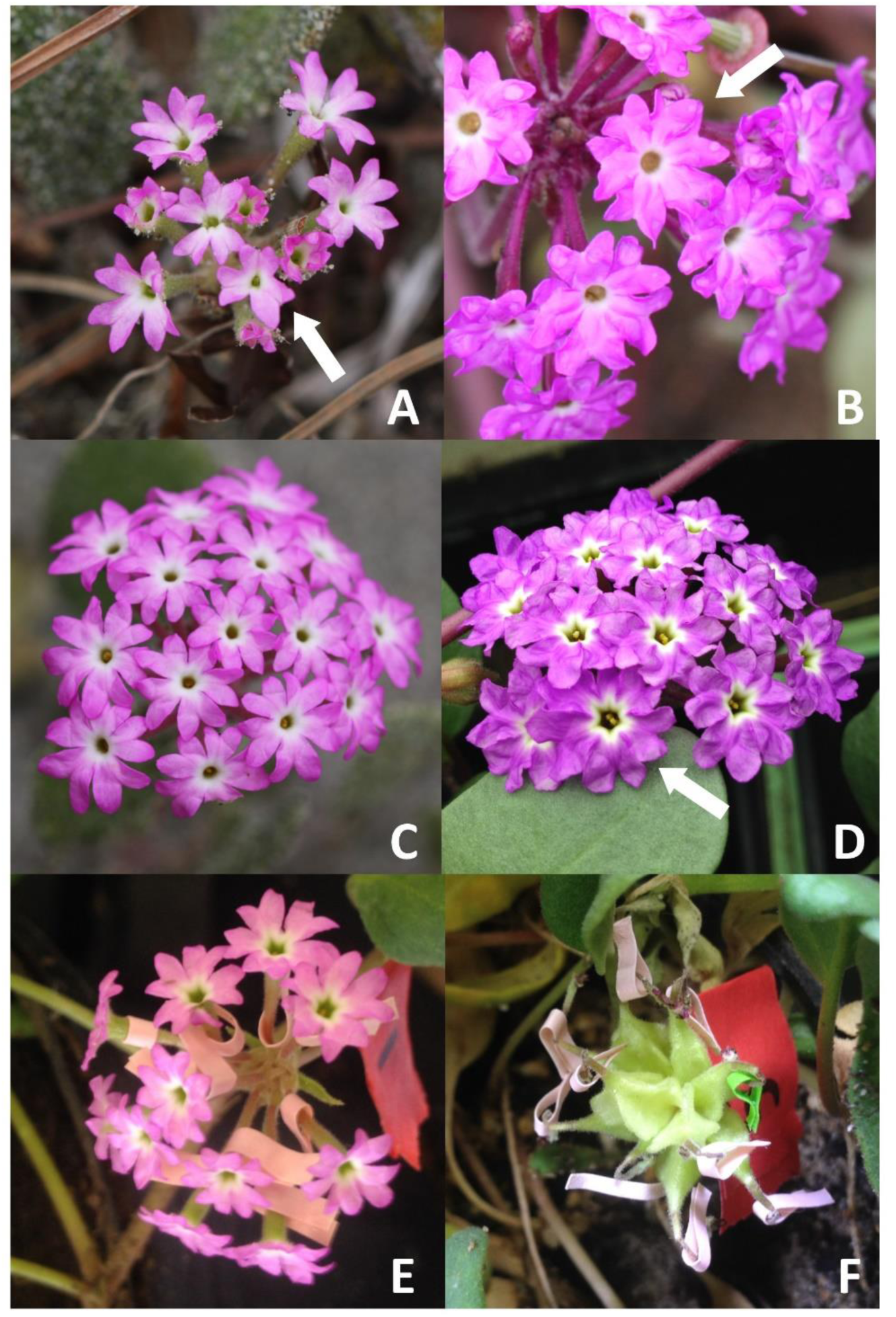
A: *Abronia umbellata* var. *breviflora* with many 4-petalled and a 3-petalled (arrow) flower. Doran Beach, Sonoma County, CA. B. *A. u.* var. *umbellata* with a 6-petalled flower (arrow). Morro Beach, San Luis Obispo County, CA. (C) *A. u.* var. *breviflora* with an atypically low number of 4-petalled flowers (none visible). Doran Beach, Sonoma County, CA. D. A. u. var. *umbellata* with a six-petalled flower (arrow). Laboratory plant. E. Individually marked flowers in autogamy experiment on *A. u.* var. *breviflora*. F. Developing fruit with tape denoting petal counts. All photos by EFL.

### Environmental or developmental effects (H5)

To determine whether petal number is influenced by environmental factors (H5), we tested for correlation between these environmental variables and each of atypical petal production (binomial response) and inflorescence mean petal number (Gaussian response) using the iNaturalist data set. Because selfing and outcrossing varieties are parapatric by latitude, we avoided confounding the effects of latitude and season with the effects of mating system by modelling petal number separately for the selfing and outcrossing portions of the species range separately using mixed-models with latitude as a continuous predictor and iNaturalist record as a random effect. To evaluate seasonal effects, we fit models with either Julian date (from Jan 1 of each year), or a squared term for Julian date (in case of changes at start and end of flowering period, e.g. McKim et al 2017), as a continuous predictor, interacting with latitude, and compared these models to null models.

### Correlation of floral size and petal numbers (H4)

To test whether flower petal number correlates with flower size, we performed analyses at the flower level, as well as at the individual and population levels. For all three levels, we used atypical petal counts, mean petals per inflorescence and floral measurements taken during the large three-population var. *breviflora* growout at Oklahoma State University. We analyzed atypical petal counts as a binomial response variable with floral size and population as fixed predictors and individual as a random effect; we used the same model structure for analyzing mean petal numbers (per inflorescence) as a Gaussian response variable. Because inner and outer flowers within inflorescences differ in size (inner are smaller in both face and tube measurements; Tillett 1967, LoPresti & Van Natto, unpublished data), we also examined whether they differed in petal numbers, as differences in meristic size altering petal number could occur within an inflorescence as well.

For a broader population level approach, we combined our iNaturalist data with field measurements of both varieties (n=407 plants, 814 measurements) taken from 26 populations across the range of *A. umbellata* by A.V.N. The flowers were chosen haphazardly from a randomly chosen branch and were not in the outer row of the inflorescence, as outer flowers are more heavily zygomorphic and more variable in measurements (Tillett 1967; Curtis 1977; Van Natto & LoPresti, pers. obs.). The average of the two measured flowers was used in all the analyses. Populations which were close by (i.e. within 0.01 degrees latitude) were combined for analyses (7 combined with nearest neighbor).

For the population level analysis of each variety, these floral size measurements were used as a predictor for atypical petal counts from the same regions (+/- 0.013 degrees; an area chosen so there would be no overlap) in the iNaturalist data set (n=453 records, 948 inflorescences, 5688 flowers) using binomial mixed models with record as a random factor. A mixed model including population floral size (from A.V.N. measurements) and Latitude (population mean) was used as a predictor for atypical flower production and mean petals per inflorescence (from iNaturalist data).

Lastly, since flowers in the outer whorl of the inflorescence are larger than inner flowers, we examined whether these size differences affected atypical floral frequency or mean petal production at the inflorescence scale in the Oklahoma State University var. *breviflora* growout. Because we had both edge and center data from the same inflorescence, we used a paired t-test to analyze each, with an arcsine transformation of the atypical flower proportion.

## RESULTS

### Hypothesis 1: Self-fertilization success of differing petal numbers

Petal number had no effect on whether *A. umbellata* var. *breviflora* flowers set seed. Overall, 76.3% of 169 4-petaled flowers set seed compared to 78.3% of 417 regular 5-petaled flowers (binomial GLM; LRT test of petal number versus an intercept-only null model χ^2^ = 0.68, df = 1, *P* = 0.41).

### Hypothesis 2: Frequency of atypical flowers across a mating system transition

The frequency of flowers with atypical petal numbers was higher for selfing than outcrossing varieties, based on both iNaturalist photographs and plants grown in a common environment. In iNaturalist photos, the frequency of atypical flowers with means taken at the inflorescence level was 20.1% among selfing var. *breviflora* records (*n* = 1526 flowers, 268 inflorescences, 120 iNaturalist records) but only 6.9% among outcrossing var. *umbellata* records (n=9899 flowers, 1540 inflorescences from 745 records) (Figs. 2, 3, 4). The varieties differed significantly in their production of atypical petals (binomial GLM; LRT test of variety versus an intercept-only null model χ^2^ = 93.6, df = 1, *P* < 0.0001; post-hoc comparison of means p < 0.0001).

**Figure 2:**
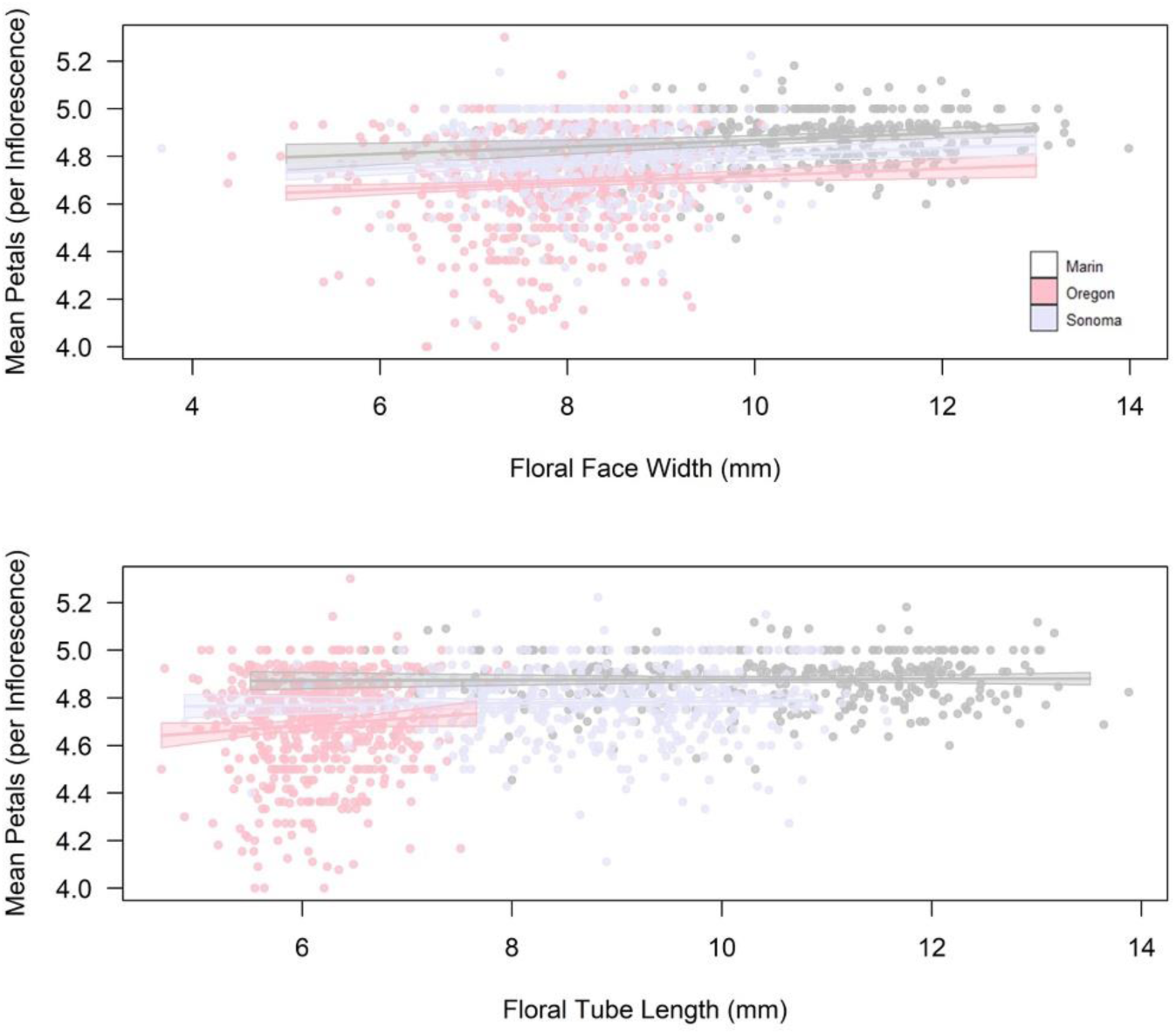
Relationship between floral size metrics and mean petal numbers in the var. breviflora growout. In the lower graph, we plotted the best-fitting regressions for each population separately; because population correlated with both mean petals and tube length, the relationship between tube length and mean petals was only significant when population was not in the model. In contrast, there was a significant additive effect of face width on mean petals; inflorescences with larger flowers also had higher mean petal number, at both the between-population and within-population level.

As predicted under correlated selection for flower size, small-flowered, selfing var. *breviflora* had a lower mean petal number than the large-flowered, outcrossing var. *umbellata* (4.80 ± 0.24 versus 4.98 ± 0.15 SD petals, respectively). The varieties differed significantly in mean petals per inflorescence (binomial GLM; LRT test of variety versus an intercept-only null model χ^2^ = 139.8, df = 1, *P* < 0.0001); confirmed with a post-hoc comparison of means. The lower mean of var. *breviflora* was due to a higher preponderance of 4-petalled flowers (19.6% of total flowers) with few 6-petalled flowers (0.3%), in contrast, var. *umbellata* had 4.6% 4-petalled and 2.1% 6-petalled.

The results from the common garden grow outs were remarkably similar in both frequency of atypical flowers and mean petals per inflorescence. In the OSU common garden, var. *breviflora* had a mean of 4.77 (± 0.18) petals per flower, with 21% 4-petalled and 0.4% 6-petaled (n=28244 flowers, 2182 inflorescences, 99 plants). In the smaller MSU common garden, var. *breviflora* had 25% four-petalled flowers (n = 656 flowers, 63 inflorescences from 10 plants) whereas var. *umbellata* had 93% 5-petalled and 7% four-petalled flowers (n= 279 flowers, 16 inflorescences, 6 plants).

### Hypothesis 2: Production of atypical flowers across three generations of inbreeding

The generation of inbreeding of var. *breviflora* did not affect production of atypical flowers at all or alter the mean petal number. A model for frequency of atypical morphs with both generation of inbreeding and population did not fit significantly better than a null with population alone (binomial GLMM; LRT χ^2^ = 1.17, df = 1, *P* =0.28), and neither was a model for mean petal number (Gaussian MM; χ^2^ = 1.27, df = 1, *P* =0.26).

### Hypothesis 3: hybridization effects on petal numbers

We found no evidence for hybridization destabilizing meristic stability; the only noteworthy result was in the opposite direction, hybridization of var. *breviflora* with *A. latifolia* reduced production of atypical flowers. For full results, see the supplementary information.

### Hypothesis 4: correlated selection on floral size

Different metrics of floral and inflorescence size correlated with both atypical flower frequency and mean petal counts (Table 3; Figure 2). These results were remarkably consistent between the data sets and varieties: as predicted, larger flowers were less likely to be atypical (i.e. a negative correlation of size and variance) and more likely to have a higher mean petal count (i.e. a positive correlation of size and mean). The full results for Table 3 are detailed in the supplementary material.

**Table 3:** Results of regressions of multiple metrics of floral size on both production of atypical petals and mean petal number. Our hypothesis of correlated selection on floral size, i.e. a decrease during the shift to autogamy in var. breviflora (Doubleday et al 2013), predicts that mean petals per flower will decrease in smaller flowers (a positive correlation) and variance will increase in smaller flowers (a negative correlation).

| Data set | Metric | Floral Face Diameter (mm) | Tube Length (mm) |
| --- | --- | --- | --- |
| Field and iNaturalist data, comparison of large-flowered var. *umbellata* to small flowered var. *breviflora* | Atypical flower production* | Negative Correlation | Negative Correlation |
|  | Mean petals/flower* | Positive Correlation | Positive Correlation |
| OSU var. *breviflora* growout, overall | Atypical flower production | Negative Correlation | Negative Correlation** |
|  | Mean petals/flower | Positive Correlation | Positive Correlation** |
| OSU var. *breviflora* growout, within inflorescence | Atypical flower production*** | Negative Correlation | Negative Correlation |
|  | Mean petals/flower*** | Positive Correlation | Positive Correlation |
| Combined A.V.N. field and iNat data, var. *breviflora* | Atypical flower production | Negative Correlation | Not measured |
|  | Mean petals/flower | Positive Correlation | Not measured |
| Combined A.V.N. field and iNat data, var. *umbellata* | Atypical flower production | No significant correlation | Not measured |
|  | Mean petals/flower | Positive correlation**** | Not measured |

\*var. *umbellata* exceeds var. *breviflora* in both of these metrics - Tillett 1967, Doubleday et al 2013, \*\*highly significant correlation disappears if population is added to model interactively or additively, \*\*\*inner flowers are smaller in both dimensions than edge flowers of the same inflorescence; this relationship, not individual measurements, was used for this analysis, \*\*\*\*591/800 records of var. *umbellata* had 5 mean petals/inflorescence, this lack of variance in response variable led to extremely bad model fit, so these results should be interpreted cautiously

**Table 4:** The results of each tested hypothesis in this study.

| Hypotheses from literature tested here: | Predictions for *Abronia umbellata* | Results |
| --- | --- | --- |
| (H1) Loss of petals increases autogamous success | In *A. u.* var. *breviflora*, fewer petalled-flowers will have higher reproductive success in greenhouse | Not supported |
| (H2) Inbreeding increases developmental instability | (1) *A. u.* var. *breviflora* (autogamous) will have increased atypical petal morphs<br>(2) Repeated inbreeding in *A. u. breviflora* will increase rates of atypical petal morphs | (1) Strongly supported, but predictions also consistent with all other hypotheses<br>(2) Not supported: no changes |
| (H3) Incompatibilities due to hybridization increase atypical petal morphs | (1) Artificially-created hybrids will have increased atypical petal morphs<br>(2) *A. u.* var. *umbellata* (xenogamous), which hybridizes with congeners commonly in the field, will have an increased atypical petal morphs in field compared to autogamous variety | (1) Not supported: hybridization stabilized petal number in var. *breviflora* x *A. latifolia* crosses<br>(2) Not supported |
| (H4) Correlated evolution with other traits (here: flower size) | (1) *A. u.* var. *breviflora* (smaller flowered) will have fewer mean petals<br>(2) Within each variety, individuals with smaller flowers will have fewer mean petals<br>(3) Within each variety, populations with smaller flowers will have fewer mean petals | (1) Strongly supported<br>(2) Strongly supported, though confounded with latitude<br>(3) Strongly supported, though confounded with latitude |
| (H5) Environmentally-driven | (1) A correlation between phenology and atypical petal morphs for each variety<br>(2) A correlation between latitude and atypical petal morphs for each variety | (1) Not supported<br>(2) Supported, though confounded with flower size |
| (H6) Ontogenetically-driven | A correlation between age of plant and atypical | Not supported, idiosyncratic relationship |

### Hypothesis 5: seasonal drivers of petal number

We found no significant effect of phenology on atypical flower production or mean petal number of either variety from the iNaturalist data set. Date (days after 1-Jan each year) had no correlation with atypical petal production either linearly or with a second-order polynomial for either variety. Mixed models including only the covariate latitude and the random effect of the record ID fit better than those containing the day for both varieties (LRT var. *breviflora*: X^2^=1.46, df=1, p=0.23; var. *umbellata*: X^2^=0.07, df=1, p=0.79). Similarly for mean petal number, a null model with only latitude fit better than a model including date for both varieties (X^2^=1.91, df=2, p=0.38 & X^2^=1.14, df=2, p=0.56).

### Hypothesis 5: environmental or genetic clines as drivers of petal number

Latitude, a proxy for either an environmental or genetic cline, predicted atypical flower production better than a null model in each variety (var. *breviflora*: X^2^=23.2, df=1, p < 0.001; var. *umbellata*: X^2^, df=1, p < 0.001). However, the amount of variance explained by latitude in these best-fitting models was small (marginal pseudo-R^2^, package *sjstats*: *breviflora* = 0.06; *umbellata* = 0.02). In both models, the random effects of record (i.e. a proxy for individual) explained somewhat more variance (conditional pseudo-R^2^, *breviflora* = 0.20, *umbellata* = 0.26). Using mean petal number instead to characterize the trend, we found mean petal number decreased with increasing latitude in both varieties. In *A. u.* var. *breviflora*, the relationship between mean petal number and latitude was stronger (marginal r^2^ = 0.13); for every increase of 1° latitude, mean petal number decreased by 0.042 (± 0.008); from the southernmost populations (~38 N) to the northernmost (~49 N), this equates to a drop in mean petal number of almost 0.5! In contrast, while the correlation between latitude and mean petal number was significant in *A. u.* var. *umbellata* as well and this model fit better than a null; the effect explained little variance (marginal r2 = 0.01) – and predicted a drop of 0.01 petals per every increase of 1° latitude (i.e. 0.05 drop across the entire range). Regardless, this result demonstrates that the direction, but not magnitude, of correlation is consistent between the varieties.

### Hypothesis 6: Ontogenic drivers of atypical petal numbers

We found a significant, yet highly idiosyncratic, effect of days since first flower on atypical flower production in var. *breviflora* (Supplementary Figure 5). In our var. *breviflora* growout, days since first flower was a significant positive predictor of atypical floral production (i.e. increasing numbers later in ontogeny). However, there was also a strong interaction of days since first flower with population, meaning that the direction of change depended on population.

## DISCUSSION

Using a combination of a large citizen-science dataset, geographic surveys, common garden experiments, and controlled hybridization and inbreeding experiments, we characterized petal number variation across the entire range of a widespread species to critically examine several hypotheses about the evolutionary causes of petal number variation in plants and meristic variation more broadly. Our data, in its totality, is most consistent with these meristic changes being a consequence of correlated evolution of smaller flowers in the selfing variety, though we cannot fully eliminate several alternative hypotheses with similar predictions (Tables 2 & 3). We were able to conclusively reject several hypotheses. We did not find support for the hypotheses of greater autogamous efficiency of reduced petal numbers (H1), short-term (up to 3 generations) inbreeding (H2:2) or hybridization (H3) leading to increased atypical petal morphs, and while evidence for environmental and ontogenetic drivers of plasticity was mixed, and the effects were too small and too population-specific to account for the overall pattern (H5/H6). We believe the approach we used can be applied to other systems to better understand this interesting, yet understudied, phenomenon.

### Evidence for correlated evolution with floral size

Floral size generally decreases substantially during transitions to selfing (Krizek and Anderson 2013), including in *A. umbellata* (Doubleday et al 2013). This transition in variety *breviflora* also correlated with a significant decrease in mean petal number and an increase in variance of petal number. These correlated results, on their own, cannot eliminate any of the remaining hypotheses, since several are agnostic to the direction of mean change (Table 1). However, given there is a strong mechanistic basis linking meristic size to organ number (e.g. Stevens et al 1972, Green 1992, Running et al 1998, Barrero et al 2006), and we supported this mechanistic relationship at every level – from within inflorescence to across the evolutionary transition to selfing – we believe it provides convincing evidence that most of the meristic variation in var. *breviflora* arises due to correlated evolution of small flowers.

Fundamentally, the two varieties of *A. umbellata* differ greatly in floral size. Given that close relatives, including *Abronia villosa*, are also obligately outcrossing and large flowered, with low rates of atypical petal production (E. LoPresti, pers. obs.), the small, often-atypical, flowers and autogamy in var. *breviflora* are both derived states (Doubleday et al., 2013). This directional relationship was strikingly obvious while going through the photographs and examining plants in the field: four-petalled morphs make up 19.4% of total flowers in var. *breviflora* (97% of all atypical morphs in this variety), but only 4.5% in var. *umbellata* (67% of atypical). Also as expected with floral size correlation, six-petalled morphs were commonly encountered in var. *umbellata* (2.1% of total, 31% of atypical), whereas they were extremely rare in var. *breviflora* (0.3% of total, 1.6% of atypical).

At a slightly finer scale, we found the same correlation. In var. *breviflora,* both floral face diameter and tube length size correlated negatively with atypical petal production and positively with mean petal number (and probably for face diameter in var. *umbellata* as well: Table 3). Finally, even within an inflorescence, smaller central flowers had higher rates of atypical petal production and lower mean petal numbers. For these reasons, we feel that a potential reduction in floral meristem size associated with smaller flowers is probably the largest contributing factor to this shift in both the production of atypical flowers (given a fairly inflexible basal state of 5 petals) and, especially, in the mean petals per flower. Given the strength of the mechanistic link between these traits in model and non-model systems (Stevens et al 1972; Green 1992; Running et al 1998, Fletcher 2001, Barrero et al 2006), we encourage comparative investigations to test the macroevolutionary implications of this so-far robust, intra-specific pattern.

### Could pollinator selection for five-petals drive this pattern?

A relaxation of pollinator-mediated selection maintaining five-petalled flower (Leppik, 1953, 1956), or simply for petal number constancy (Herrera, 2009) given the basal state of 5 petals, during the evolution of autogamy in var. *breviflora* hypothetically could explain this transition. Pollinator-mediated selection on petal number might require pollinators to be able to count petals, which bees have been found to be capable of (Leppik, 1953, 1956; Lehrer et al., 1995) but Lepidoptera have not (Leppik, 1954, 1955). *Abronia umbellata* is likely primarily moth pollinated; Doubleday and Eckert (2018) found it had higher floral visitation during the day, but all but a very small amount of pollination occurred at night. Given these lines of evidence, it seems highly unlikely that the lower production of atypical morphs in var. *umbellata* is due to pollinator-mediated selection (especially given that 5-petalled flowers as an ancestral state predate Nyctaginaceae and occurs despite diverse pollinators across the Carophyllales). Correlated selection on numbers of other meristic traits by pollinators is possible, though implausible. *Abronia* has one perianth whorl and the stamens and the single carpel are both contained inside the tube and difficult to see, or count. While the predictions of these hypotheses are largely consistent with our results (Tables 2 & 3), we find them far less likely to explain the meristic changes observed than correlated evolution with floral size.

### The effect of inbreeding

While our short-term, three-generation, laboratory inbreeding had no effect on atypical flower production in *A. u. breviflora*, it is still possible that long-term inbreeding may be partially responsible for the observed high rates of atypical petal production in these selfers. In contrast to the self-incompatible *A. u. umbellata*, *A. u.* var. *breviflora* reproduces almost solely autogamously (Doubleday et al., 2013; Van Natto, 2020), and therefore, individuals in our study were probably inbred prior to beginning those lab lines. Consistent with this interpretation, Greer (2016) and Van Natto (2020) found that there was little genetic variation in *A. u.* var. *breviflora*. Inbreeding may increase aberrant morphs in meristic traits (Sherry and Lord, 1996), and given a lack of xenogamous pollination and that atypical flowers set seed autogamously at the same rate as typical flowers, these mutants potentially carry little cost in these populations. While this is still a viable hypothesis in our system, and may explain some variance unexplained simply by reduction in floral size, the only other evaluation of this hypothesis that we know of, which used data from Schlichting and Levin (1986) on xenogamous *Phlox drummondii*, found that inbreeding decreased the proportion of atypically petalled flowers, rather than increasing it (Mickley and Schlichting, unpublished data). While our short-term test could not conclusively eliminate this hypothesis, it certainly did not support it, either.

Ultimately, any broad conclusions would need to characterize atypical floral production across multiple transitions. While the selfing *A. u. breviflora* had reduced mean petal numbers, which is consistent with several hypotheses about atypical petal production; it also correlated with a reduction in floral size during the same transition. In large, phylogenetic investigations across multiple transitions, we hypothesize that selection for smaller flowers in selfing lineages will produce a directional loss in petals, whereas any relaxation of pollinator constancy would increase variance, but could be in a gain, or a symmetric increase around the mean (i.e. variance increases independent of the mean, which may be unchanged).

### No effect of petal number on autogamy

The relationship between small-flowered selfing species and loss of petals led Monnaiux et al (2016) to propose an adaptive hypothesis. They reasoned that petal expansion assists in opening of the floral bud, and therefore a reduction in petal number might delay bud opening and increase successful autogamous fertilization. In our laboratory study, we found no support for increased autogamous success of four-petalled morphs compared to five-petalled morphs in the autogamous *A. u.* var. *breviflora*. There is a possibility that in the field, four-petalled morphs may get fewer, shorter, or less effective pollinator visits and thus, may have lower pollen removal and therefore higher autogamous success; our laboratory study could not examine that hypothesis. Additionally, we attempted and failed to find any distinguishing characteristics in fruit resulting from flowers of differing petal numbers, meaning a *post hoc* analysis could not be done from field seed collections as we had initially hoped. Any test for autogamy in a field setting would therefore require marking individual flowers, as we did in the laboratory. Nevertheless, our study was the first to test this interesting hypothesis and we rejected it; however, our methods are easily replicated in other systems to obtain a stronger, broader result.

### Population, not phenology, contributed strongly

In other systems, changes in petal numbers occur plastically in response to environmental variables (e.g. Huether, 1968; McKim et al., 2017), the two which we could test with the large iNaturalist data set were latitude and phenology (both as proxies for climatic variables or day length). We did not find any effect of phenology on either variety in our iNaturalist data set. Coupled with the consistent patterns of atypical petal production and mean petal number between field and common garden plants, the lack of a seasonal correlation is more evidence that the observed production of petal morphs is largely heritable and the latitudinal effect should be investigated to parse out population differentiation across latitudes or a plastic response to environmental differences across latitudes.

In both varieties, though to a far lesser extent in var. *umbellata*, latitude was correlated with atypical petal number; as latitude increased mean petal number decreased. Populations along this gradient strongly differed in mean petal number (especially in var. *breviflora*, Table 2), as well as in floral size. While this effect could have been driven by environmental factors associated with latitude, we feel that is unlikely. Instead, the linear nature of this species range and the fragmentary population structure of the northern populations make it likely that this gradient, too, is due to a genetic cline underlying both floral size and meristic stability. The smaller populations of var *breviflora* in the northern part of their range were homozygous at every allele and had no private alleles (Van Natto and Eckert 2022). Those northern populations also had the smallest flowers, and were the only ones where we noted appreciable numbers of three-petalled flowers.

### No consistent effect of hybridization on petal number variation

Hybrids of *A. umbellata* did not exhibit increases in atypical petal number, in fact, hybrids of var. *breviflora* had reduced atypical petal numbers, in direct contrast to our hypothesis. Additionally, since var. *umbellata* hybridizes often and var. *breviflora* does not (Tillett 1967; Van Natto 2020), we can, on its face, entirely reject the possibility that introgression is responsible for the derived condition of increased atypical petal production in var. *breviflora*. Alternately, could a legacy of introgression reduce petal number variation in *A. u.* var. *umbellata*? Hybridization had no significant effect on atypical petal production in var. *umbellata*, but stable petal numbers of five are the norm in *Abronia* (LoPresti, pers. obs.), and therefore, meristic instability is a derived condition of var. *breviflora* instead of a basal character of the clade. Our data on greenhouse hybrids (Supplementary material) shows that most hybrids of var. *umbellata* we made had low production of atypical flowers. Demonstrating that the low levels of introgression into this variety (Tillett, 1967; Van Natto and Eckert 2020) stabilizes petal number in natural populations would require far more investigation than this survey study could provide, and while not a possibility that our data suggest is at all likely, we cannot conclusively reject it either.

### Conclusions

Integrating a common garden study, crossing experiment, and large-scale field citizen science data, we characterized an intriguing shift in atypical petal number during a transition from outcrossing to autogamy in a coastal sand verbena. The increase of atypical petal number of the selfing variety was not consistent with directional selection caused by increased autogamous success, nor was it driven by seasonal environmental changes, as in other similar systems. Our data most strongly suggested that correlated selection decreasing flower size during a transition to selfing drove the reduction of mean petals, a result predicted by studies on floral initiation. We also demonstrate the power and ease of using high-quality photographs from citizen-science efforts to score easily quantifiable traits across range and seasons to generate a data set which would have been logistically difficult to get with fieldwork or herbarium specimens.

## ACKNOWLEDGEMENTS

The authors thank the many iNaturalist contributors for detailed photographs and identifications; the ‘data’ here was gathered by hundreds of people with cameras! We thank past and present members of the Weber lab, especially Caroline Edwards and Carina Baskett, for comments on the manuscript and on the project overall. Laura Doubleday gave us invaluable information on *Abronia umbellata*. Rick Karban, Patrick Grof-Tisza, Dena Grossenbacher, and Kimiora Ward accompanied and assisted EL on fieldwork and seed collecting of *Abronia umbellata* during 2015-2018 seasons and Carolyn Graham, Bruce Martin, and Madi Stessman assisted in caring for the common garden growouts. AD, MF, and CG were funded by start-up funds from Oklahoma State University. EFL was funded by NSF-PRFB #1708942, MGW was funded by NSF-DEB #1831164. CGE and AVN were funded by a Discovery Grant from the Natural Sciences and Engineering Research Council of Canada (NSERC). The lead author thanks all coauthors for putting up with the glacial pace that this project, begun in 2018, and the manuscript, with 33 drafts exchanged, progressed at.

## DATA AVAILABILITY

Data and code are archived on FigShare with DOIs listed in the reference section (note: I don’t see exactly how to do this in the article guidelines on the AJB website, so I just took the FigShare suggested format). The data sets are: (1) the iNaturalist data (including pure varieties and interspecific hybrids), (2) the autogamy lab data, (3) field population data from A.V.N (4) the OSU growout var. *breviflora* data, (5) collection locations and herbarium specimen numbers for plants used in the common garden studies. Lastly, there are a series of three R scripts uses the attached data sheets for the analyses in the paper. The limited hybridization data from the greenhouse is presented in the supplementary table. Much more data on hybridization (including more crosses of *A. umbellata*) will be available when the many-year, whole-genus study from which those data were taken is completed.

## ONLINE SUPPORTING INFORMATION

Additional Supporting Information may be found online in the supporting information section at the end of the article.

## Supplementary information

### Detailed results for correlated selection on floral size; i.e. Table 3

#### OSU var. breviflora growout, overall

Floral face diameter was a significant predictor of both mean petal number and atypical floral production; it also varied by population (Main text figure 2; Supplementary Figure 1). A Gaussian mixed-model for mean petal number with individual as a random effect included both floral face diameter and population additively as predictor variables; adding an interactive term did not improve fit (X^2^ = 1.17, df=2, p =0.56), though the additive model fit significantly better than a population-only null model (X^2^ = 8.14, df=1, p =0.004). A binomial mixed model for atypical petal production with individual as a random effect included both floral face diameter and population additively as predictor variables; adding an interactive term did not improve fit (X^2^ = 1.70, df=2, p =0.43), though the additive model fit significantly better than a population-only null model (X^2^ = 6.65, df=1, p = 0.010). These results demonstrate that both within, and among, populations, floral face diameter correlated strongly with mean petal number and atypical petal production in the hypothesized directions.

Floral tube length also varied by population (Supplementary Figure 2). The same modelling approach was conducted as with floral face diameter; however, models with population included did not improve with the addition of tube length. If population was removed from the model, tube length was an extremely significant predictor, improving fit over intercept-only null models for both mean petal number (X^2^ = 33.99, df=1, p < 0.0001), as well as atypical petal production (X^2^ = 40.86, df=1, p < 0.0001). These results suggest that floral tube length correlated strongly between populations with mean petal number and atypical petal production, but this effect was not seen within populations.

#### OSU var. breviflora growout, within inflorescence

On the within inflorescence scale, while not as pronounced as the ray and disc flowers of asters, *Abronia* have a similar arrangement of flowers in the inflorescence with larger, more heavily zygomorphic outer flowers and smaller, nearly actinomorphic inner flowers. The inner flowers of all species are consistently smaller in every dimension, including both floral face and tube length, and produce smaller seeds (LoPresti, unpublished data). Inner flowers had significantly lower mean petal number compared to the outer flowers on the same inflorescences (Supplementary Figure 3, paired t-test, t=17.175, df = 2177, p<0.0001). These smaller inner flowers also had ~8% (observed) higher rates of atypical petal production at the inflorescence level in the var. *breviflora* growout (Supplementary Figure 4, paired t-test of arcsin-transformed proportions: t=-7.99, df = 2177, p<0.0001).).

#### Combined Alyson Van Natto field and iNat data, var. breviflora

Both mean petal number and atypical petal production were significantly predicted by floral face in this dataset (floral tube was not measured). A model for mean petals, with both floral face diameter and latitude as fixed effects, with iNaturalist record number as a random effect, fit significantly better than one including only latitude (X^2^ = 11.78, df=1, p = 0.0006). The same result was found with a binomial model with the same structure for atypical petal production (X^2^ = 4.05, df=1, p = 0.044).

#### Combined Alyson Van Natto field and iNat data, var. umbellata

A model for mean petals, with both floral face diameter and latitude as fixed effects, with iNaturalist record number as a random effect, fit significantly better than one including only latitude (X^2^ = 5.85, df=1, p = 0.016). However, the model diagnostics were very poor, due to lack of variance in the response variable (which transforming cannot fit). Almost 75% (591/800) of the inflorescences of var. *umbellata* in this data set had a mean petal number of 5. Therefore, these results should be taken as suggestive of a pattern, but not absolutely conclusive. Floral face did not significantly improve the fit of a latitude-only null model with record as a random effect for atypical floral production (X^2^ = 0.33, df=1, p = 0.57).

### Hybridization results (H3)

Morphological hybrids of var. *umbellata* did not significantly differ from the pure variety in production of atypical flowers (6.9% in pure var. *umbellata*, 4.7% hybrids with *latifolia* and 12.5% hybrids with *maritima*). A model with the hybrids as factors did not fit better than a null model without (LRT: χ^2^=2.72, df=2, p = 0.26). Mean petals similarly did not significantly differ (4.97, 5.02, 5.05, respectively) and adding hybrid did not improve fit over a null model (LRT: χ^2^=3.39, df=2, p = 0.18). In contrast to the previous results and to our hypothesis, hybrids of var. *breviflora* with *A. latifolia* had reduced production of atypical petals from the pure species (3% versus 20%); a marginally significant difference (coefficient of hybridization −2.10 + 1.17; z = 1.78, p=0.07), though this model fit significantly better than a null without the hybridization factor (χ^2^=4.45, df=1, p = 0.03). Results from the limited sample of lab crosses were idiosyncratic; some had elevated, some had reduced variation in atypical flowers (Supplementary Table 1).

---

**Supplementary Figure 1:**
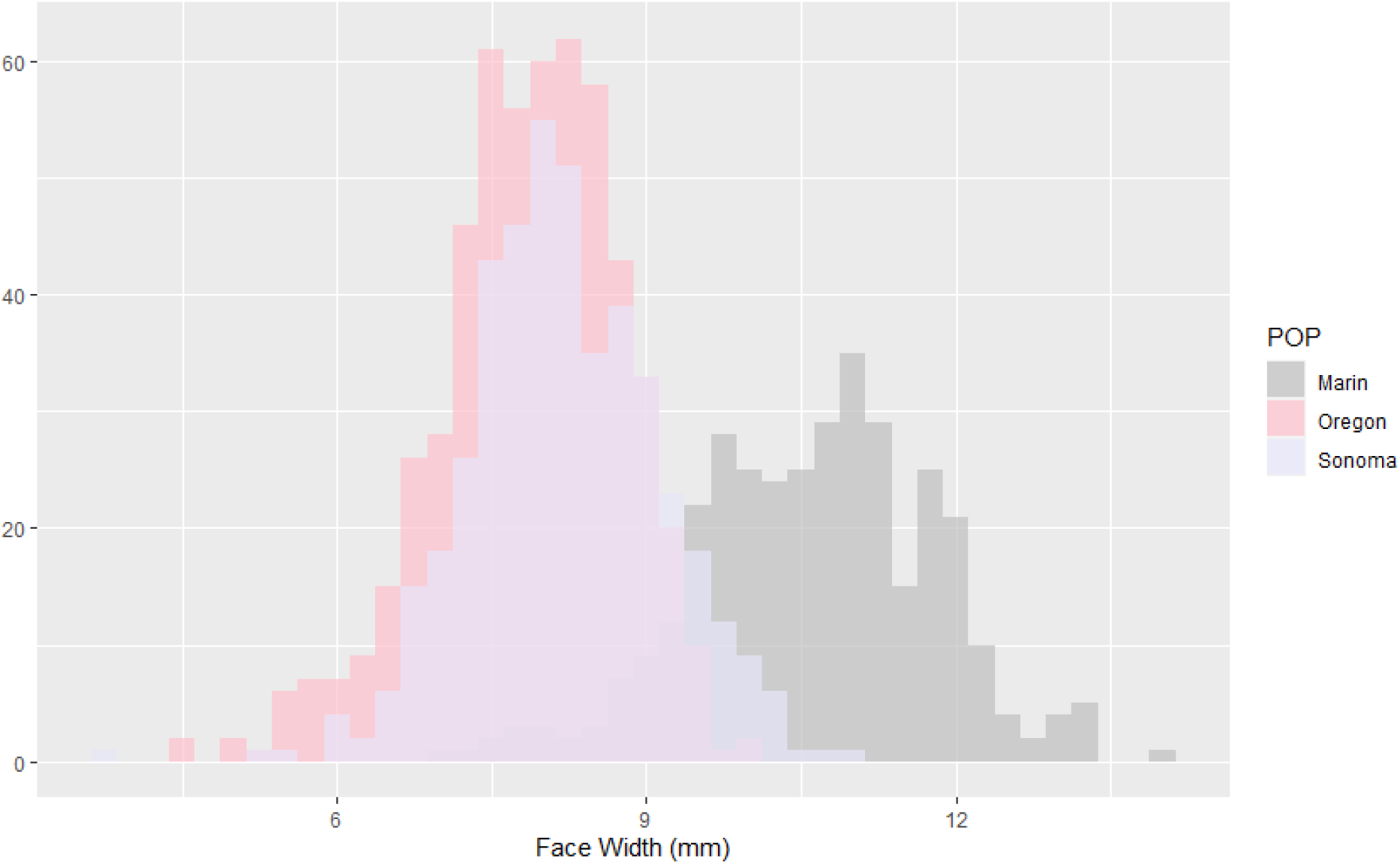
Mean floral face width (per inflorescence, most with one, some with two flowers measured) of var. *breviflora*, by population.

**Supplementary Figure 2:**
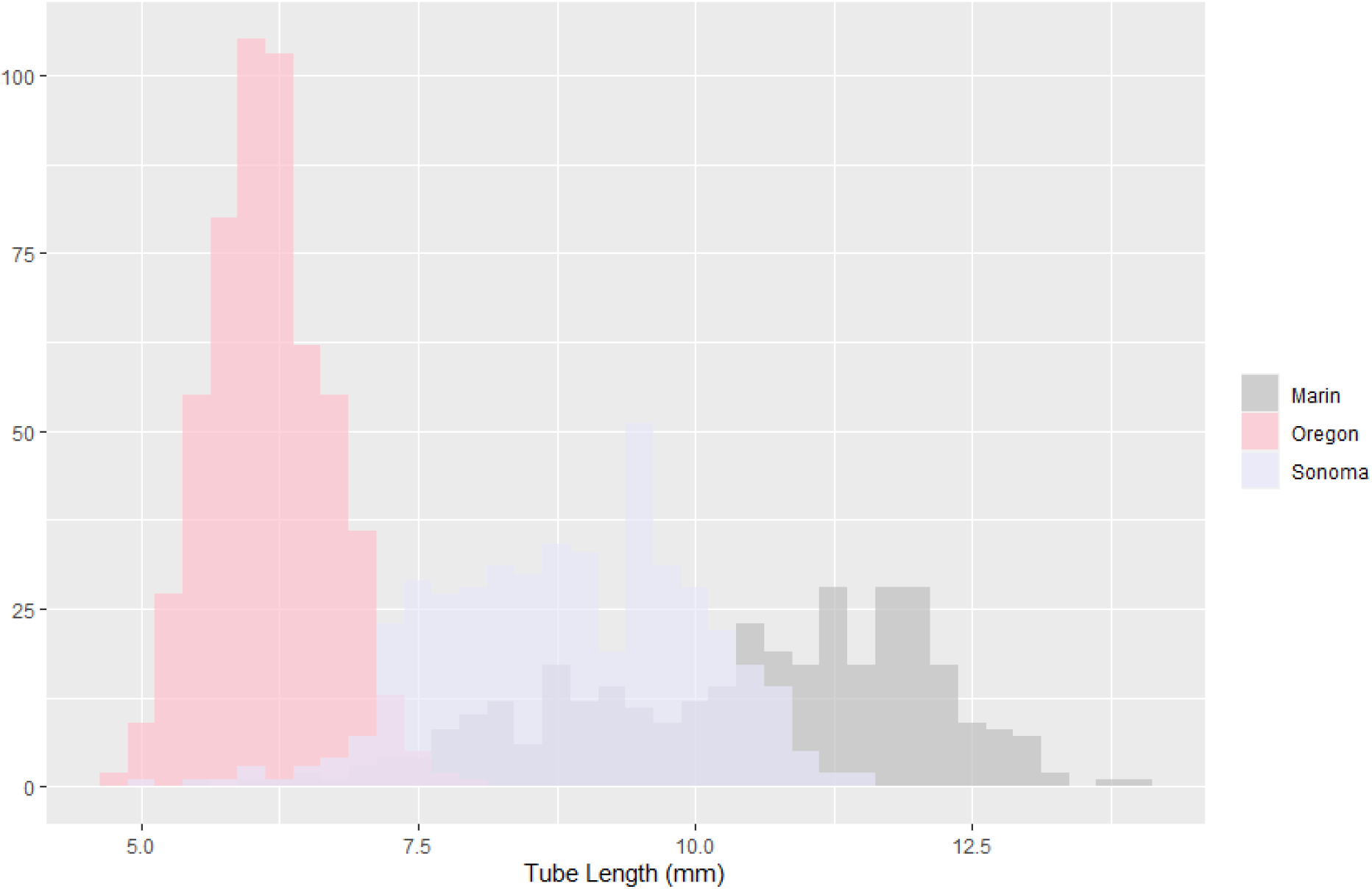
Mean floral tube length (per inflorescence, most with one, some with two flowers measured) of var. *breviflora*, by population.

**Supplementary Figure 3:**
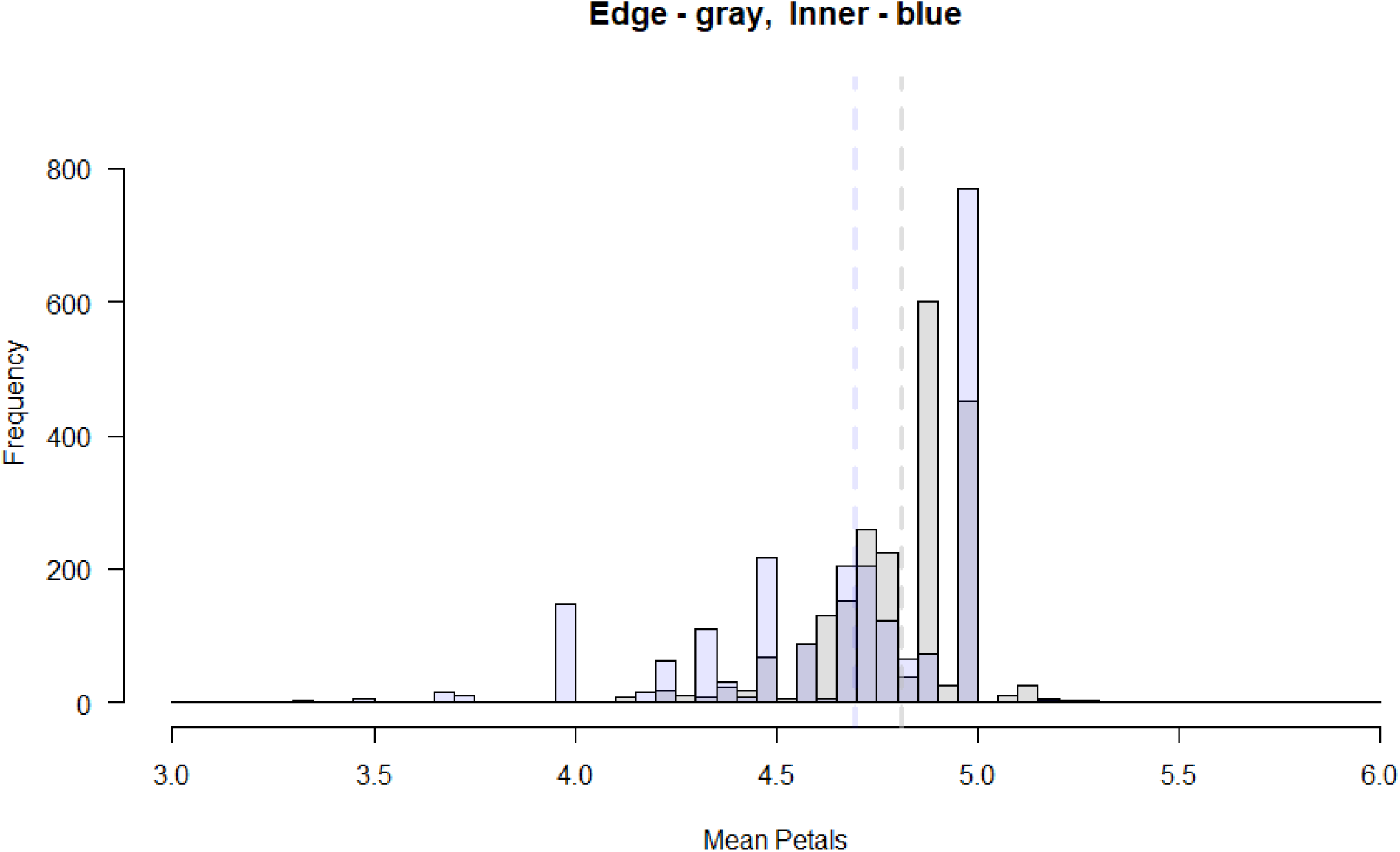
Mean petal number, per inflorescence, by flower position within the inflorescence (populations not significant factors). Means denoted by dotted lines.

**Supplementary Figure 4:**
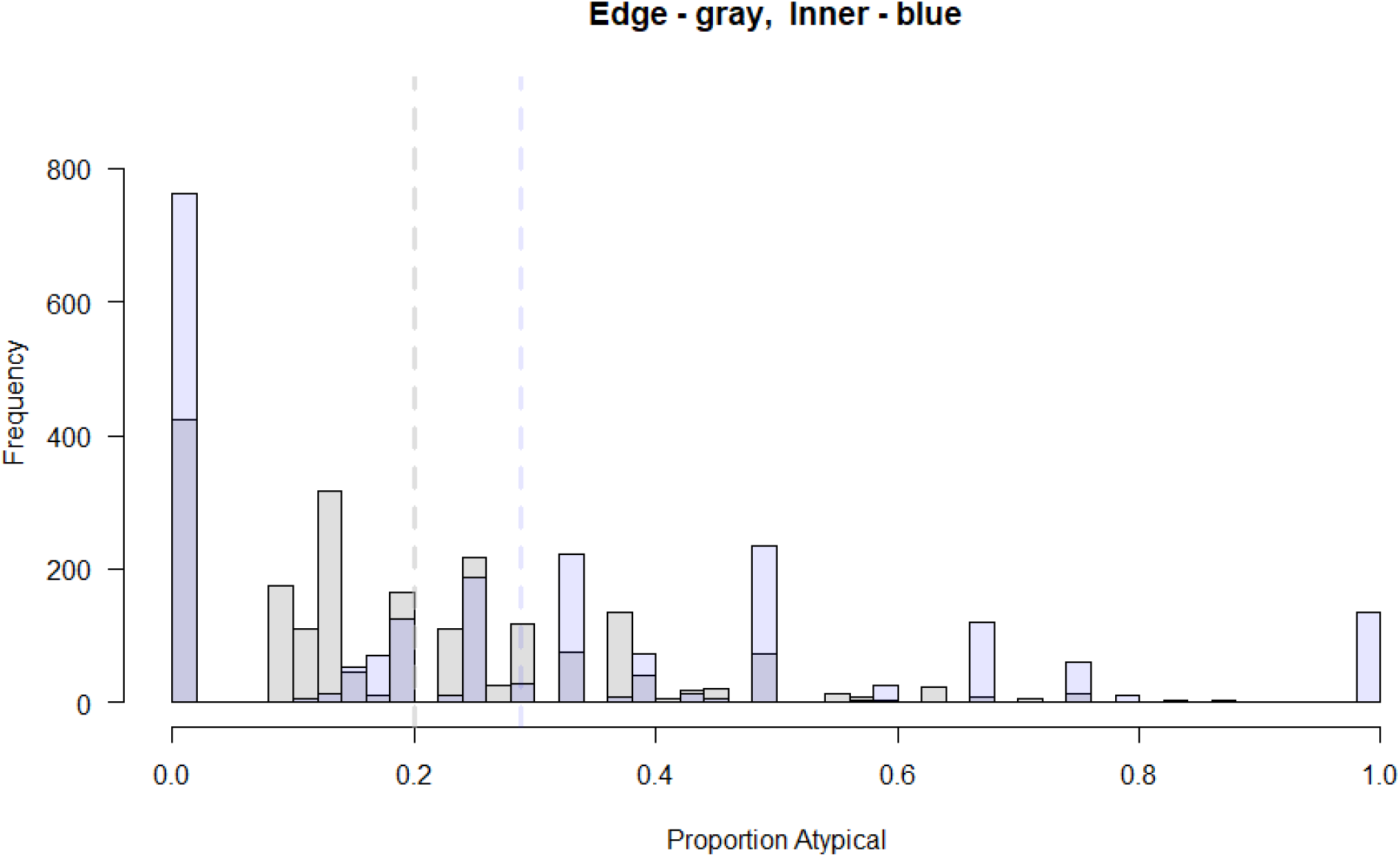
Proportion atypical petals, by flower position within the inflorescence (populations not significant factors). Means denoted by dotted lines.

**Supplementary Figure 5:**
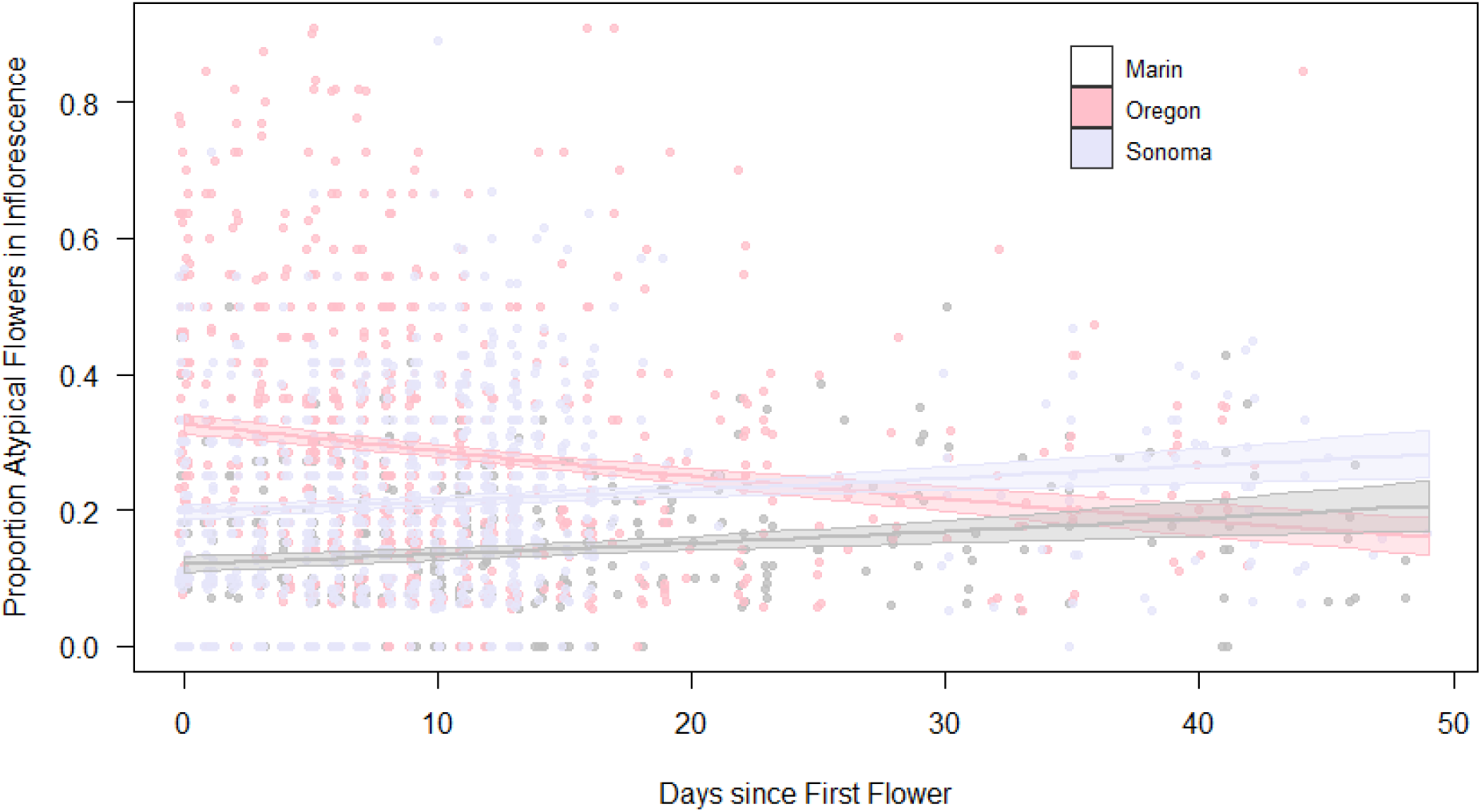
The effect of days since first flower (dsff) on atypical petal production. Predictions are from a binomial model with population*dsff;

**Supplementary Figure 6:**
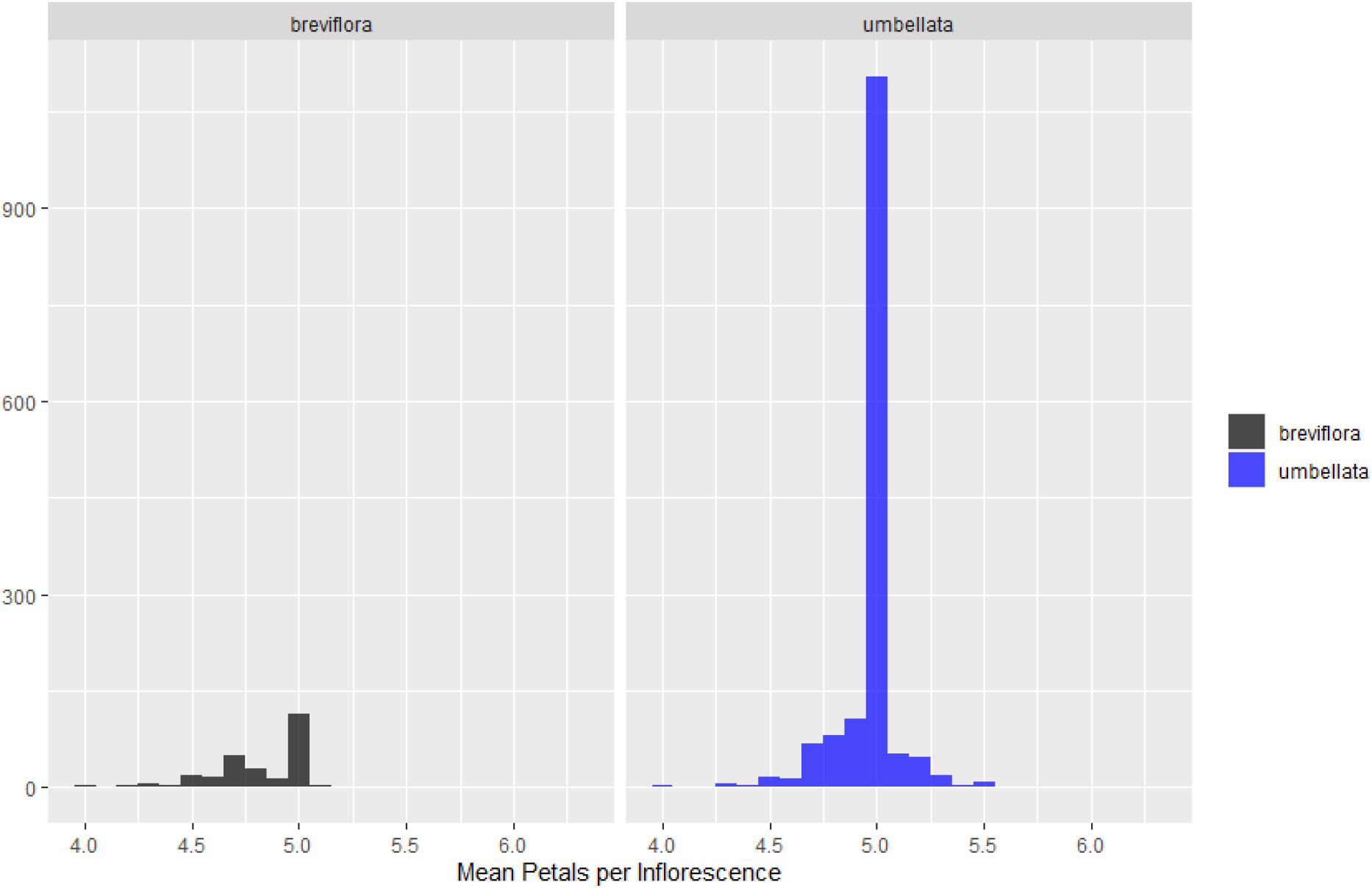
Histogram of mean petal number, by variety, from the iNaturalist data.

**Supplementary Figure 7:**
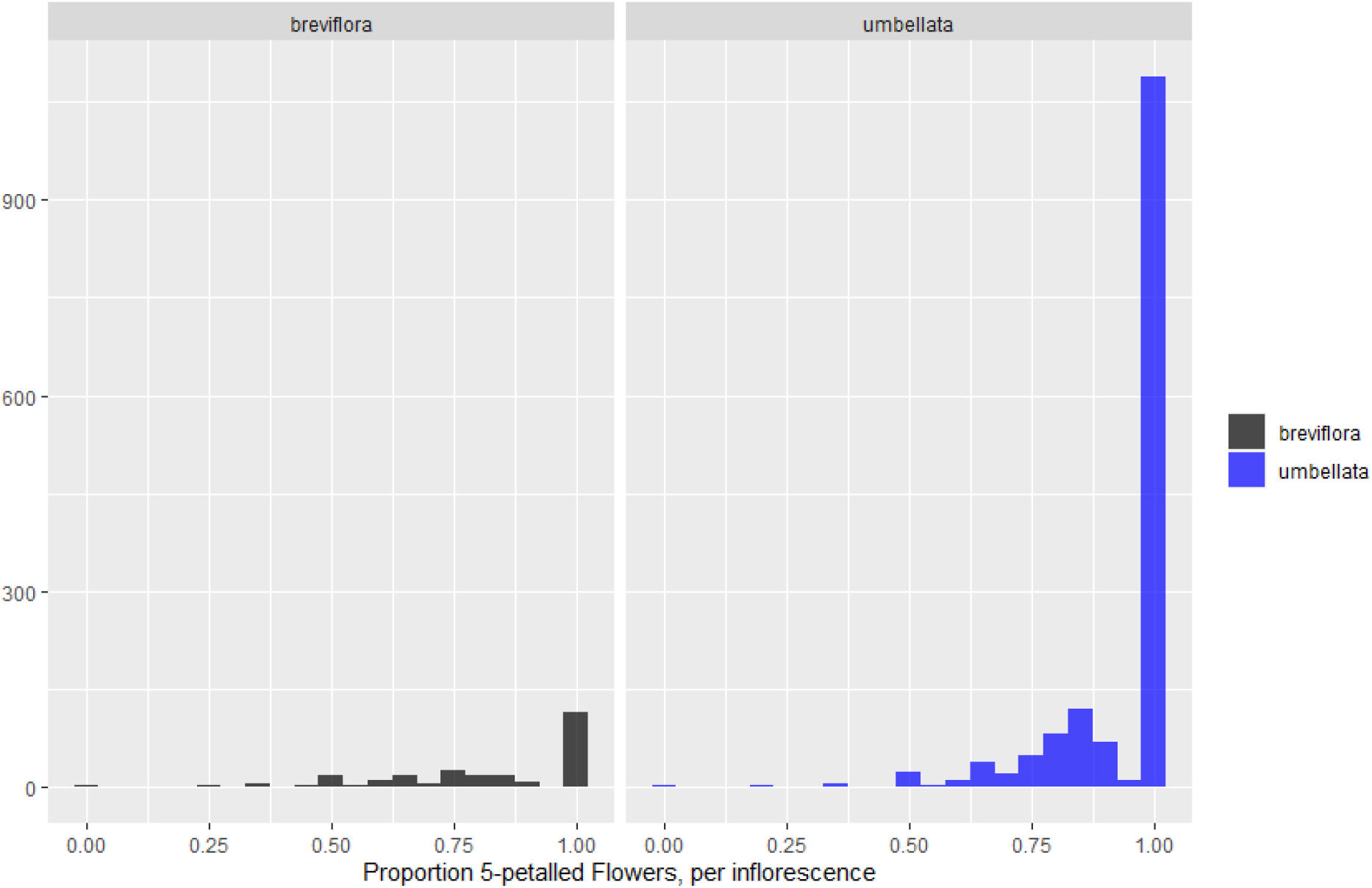
Histogram of proportion 5-petalled flowers, by variety, from the iNaturalist data.

## Notes

https://figshare.com/projects/Abronia_umbellata_petal_number/156777

### Competing Interest Statement

The authors have declared no competing interest.

### Summary of Updates

A lot of new data from a variety of people (field data from AVN and CGE; photo analysis from CG; a big growout from AD, MF, SJ, KT & EFL). The conclusions remain almost unchanged, though we now have a bit more confidence due to all the different data sets now incorporated!

## LITERATURE CITED

Agren, J., and D.W. Schemske. 1995. Sex Allocation in the Monoecious Herb Begonia semiovata. Evolution 49: 121–130.

Alpi, A., M. Buiatti, and S. Baroncelli. 1968. Some data on the polygenic control of two quantitative traits in a vegetatively propagated flower plant, the carnation. Theoretical and Applied Genetics 38: 298–300.

Bachmann, K., K. Chambers, and H. Price. 1981. Genetic determination of pappus part number in the annual hybrid Microseris B 87 (Asteraceae - Lactuceae). Plant Systematics and Evolution 138: 235–246.

Bammi, R.K., and H.P. Olmo. 1966. Cytogenetics of *Rubus*. V. Natural Hybridization Between *R. procerus* P. J. Muell. and *R. laciniatus* Willd. Evolution 20: 617–633.

Barrero, L.S., B. Cong, F. Wu, and S.D. Tanksley. 2006. Developmental characterization of the fasciated locus and mapping of Arabidopsis candidate genes involved in the control of floral meristem size and carpel number in tomato. Genome 49: 991–1006.

Bletsos, F.A., D.G. Roupakias, M.L. Tsaktsira, A.B. Scaltsoyjannes, and C.C. Thanassoulopoulos. 1998. Interspecific hybrids between three eggplant (Solanum melongena L.) cultivars and two wild species (Solanum torvum Sw. and Solanum sisymbriifolium Lam.). Plant Breeding 164: 159–164.

Byerley, M. 2006. Patterns and consequences of floral formula variation in Phlox (Polemoniaceae). PhD Thesis. Colorado State University.

Cantino, P.D., J.A. Doyle, S.W. Graham, W.S. Judd, R.G. Olmstead, D.E. Soltis, P.S. Soltis, and M.J. Donoghue. 2007. Towards a phylogenetic nomenclature of Tracheophyta. Taxon.

Choi, K., J.S. Kim, and J.H. Pak. 2001. Natural hybridization between Pseudostellaria davidii and Pseudostellaria palibiniana (Caryophyllaceae). Plant Species Biology.

Delesalle, V.A., and S.J. Mazer. 2014. The structure of phenotypic variation in gender and floral traits within and among populations of Spergularia marina (Caryophyllaceae). American Journal of Botany 82: 798–810.

Doubleday, L.A.D., and C.G. Eckert. 2018. Experimental evidence for predominant nocturnal pollination despite more frequent diurnal visitation in Abronia umbellata (Nyctaginaceae). Journal of Pollination Ecology 22: 67–74.

Doubleday, L.A.D., R.A. Raguso, and C.G. Eckert. 2013. Dramatic vestigialization of floral fragrance across a transition from outcrossing to selfing in Abronia umbellata (Nyctaginaceae). American Journal of Botany 100: 2280–2292.

Douglas, N.A. 2008. Tripterocalyx carneus (Nyctaginaceae) Is Self-Compatible. The Southwestern Naturalist 53: 403–406.

Ellstrand, N.C. 1983. Floral Formula Inconstancy Within and Among Plants and Populations of Ipomopsis aggregata. Botanical Gazette 144: 119–123.

Endress, P.K., 2001. Origins of flower morphology. Journal of Experimental Zoology 291: 105–115.

Fletcher, J.C. 2001. The ULTRAPETALA gene controls shoot and floral meristem size in Arabidopsis. 128: 1323–1333.

Galen, C. 1999. Why Do Flowers Vary? BioScience 49: 631.

Golding, Y.C., M.S. Sullivan, and J.P. Sutherland. 1999. Visits to manipulated flowers by Episyrphus balteatus (Diptera: Syrphidae): partitioning the signals of petals and anthers. Journal of Insect Behavior 12: 39–45.

Grant, V. 1956. The genetic structure of races and species in Gilia. In Advances in Genetics, 55–87. Academic Press.

Greer, S.U. 2006. Genomic consequences of mating system evolution in the Pacific coastal dune endemic Abronia umbellata (Nyctaginaceae). Queens University, Kingston, Ontario, Canada.

Herrera, C.M. 2009. Multiplicity in Unity. Chicago, U. University of Chicago Press.

Hoffmann, A.A., P.C. Griffin, and R.D. MacRaild. 2009. Morphological variation and floral abnormalities in a trigger plant across a narrow altitudinal gradient. Austral Ecology 34: 780–792.

Huether, C.A. 1968. Exposure of natural genetic variability underlying the penamerous corolla constancy in Linanthus androsaceus ssp androsaceus. Genetics 60: 123–146.

Huether, C.A. 1969. Constancy of the pentamerous corolla phenotype in natural populations of Linanthus. Evolution 23: 572–588.

Katsuyoshi, K., and J. Harding. 2015. Genetic and environmental variation for corolla traits in Portulaca grandiflora. Journal of Horticultural Science 44: 37–47.

Keeler, K.H. 1981. Function of Mentzelia nuda (Loasacae) postfloral nectaries in seed defense. American Journal of Botany 68: 295–299.

Keeler, K.H. 1979. Nocturnal pollination of Abronia fragrans (Nyctaginaceae). Southwestern Naturalist 24: 692–693.

Kido, A., and M. Hood. 2020. Mining new sources of natural history observations for disease interactions. American Journal of Botany 107: 3–11.

Kitazawa, M.S., and K. Fujimoto. 2014. A developmental basis for stochasticity in floral organ numbers. Frontiers in Plant Science 5: 1–14.

Lambrecht, S.C. 2013. Floral water costs and size variation in the highly selfing Leptosiphon bicolor (Polemoniaceae). International Journal of Plant Sciences.

Lehmann, N.L., 1987. Floral formula variation in Phlox drummondii Hook. University of Texas at Austin. Master’s Thesis.

Leppik, E.E., 1953. The Ability of Insects to Distinguish Number. American Naturalist 87: 229–236.

Levy. 1997. Nonhomeotic meristic flower mutants in *Phacelia dubia*. Journal of Heredity: 88:31–37

LoPresti, E., 2023. OSU breviflora data. figshare. Dataset. https://doi.org/10.6084/m9.figshare.21817668.v1

LoPresti, E., 2023. iNaturalist data matching AVN measured populations. figshare. Dataset. https://doi.org/10.6084/m9.figshare.21817680.v1

LoPresti, E., A Van Natto. 2023. Alyson Van Natto Petal Measurements. figshare. Dataset. https://doi.org/10.6084/m9.figshare.21817695.v1

LoPresti, E., 2023. iNaturalist Data. figshare. Dataset. https://doi.org/10.6084/m9.figshare.21817716.v1

LoPresti, E., 2023. Hybrid Abronia Data. figshare. Dataset. https://doi.org/10.6084/m9.figshare.21817725.v1

LoPresti, E. and C. Edwards, 2023: R Script for iNaturalist Data. figshare. Software. https://doi.org/10.6084/m9.figshare.21817665.v1

LoPresti, Eric and M. Foisy, 2023. Two scripts for the cleaning and analysis of the OSU breviflora data. figshare. Software. https://doi.org/10.6084/m9.figshare.21817671.v1

Lowndes, A.G., 1931. Note on Individual Variation in Paris quadrifolia L. New Phytologist 30: 298–299.

Mathiasen, R.L. 1982. Taxonomic studies of dwarf mistletoes (Arceuthobium ssp.) parasitizing Pinus strobiformis. Great Basin Naturalist 42: 120–127.

Mazer, S.J., V.A. Delesalle, and P.R. Neal. 1999. Responses of Floral Traits to Selection on Primary Sexual Investment in Spergularia marina: The Battle between the Sexes. Evolution 53: 717–731.

McKim, S.M., A.L. Routier-Kierzkowska, M. Monniaux, D. Kierzkowski, B. Pieper, R.S. Smith, M. Tsiantis, and A. Hay. 2017. Seasonal regulation of petal number. Plant Physiology.

Mickley, J., 2017. The Adaptive Nature of Stasis for Petal Number: Can Pollinator-Mediated Stabilizing Selection Explain Five-petaled Flowers? Storrs: University of Connecticut.

Mickley, J., and C. Schlichting. 2018. Revisiting an old question in California botany : Why do many plant species have five-petaled flowers? Mojave National Preserve Science Newsletter 2018: 13–16.

Monniaux, M., B. Pieper, and A. Hay. 2016. Stochastic variation in Cardamine hirsuta petal number. Annals of Botany.

Monniaux, M., B. Pieper, S.M. McKim, A.L. Routier-Kierzkowska, D. Kierzkowski, R.S. Smith, and A. Hay. 2018. The role of APETALA1 in petal number robustness. eLife.

Pélabon, C., M.L. Carlson, T.F. Hansen, N.G. Yoccoz, and W.S. Armbruster. 2004. Consequences of inter-population crosses on developmental stability and canalization of floral traits in Dalechampia scandens (Euphorbiaceae). Journal of Evolutionary Biology. 17: 19–32.

Pieper, B., M. Monniaux, and A. Hay. 2016. The genetic architecture of petal number in Cardamine hirsuta. New Phytologist 209: 395–406.

Roman, H., M. Rapicault, A.S. Miclot, M. Larenaudie, K. Kawamura, T. Thouroude, A. Chastellier, et al. 2015. Genetic analysis of the flowering date and number of petals in rose. Tree Genetics and Genomes 11:. Available at: http://dx.doi.org/10.1007/s11295-015-0906-6.

Roy, S. 1963. The variation of organs of individual plants. Journal of Genetics 58: 147–176.

Running, M. P., J. C. Fletcher, and E. M. Meyerowitz. 1998. The WIGGUM gene is required for proper regulation of floral meristem size in Arabidopsis. Development 125: 2545–2553.

Saunders, E.R. 1934. A study of Veronica from the viewpoint of certain floral characters. Journal of the Linnean Society of London, Botany 49: 453–493.

Schemske, D.W. 1978. Sexual reproduction in an Illinois population of Sanguinaria canadensis L. The American Midland Naturalist 100: 261–268.

Schlichting, C. D., & Levin, D. A. 1986. Phenotypic plasticity: an evolving plant character. Biological Journal of the Linnean Society, 29: 37–47.

Shepherd, K.A., T.D. Macfarlane, and M. Waycott. 2005. Phylogenetic analysis of the Australian Salicornioideae (Chenopodiaceae) based on morphology and nuclear DNA. Australian Systematic Botany 18: 89–115.

Sherry, R.A. and Lord, E.M., 1996. Developmental stability in flowers of Clarkia tembloriensis (Onagraceae). Journal of Evolutionary Biology, 9(6), pp.911–930.

Soltis, P.S. & Soltis, D.E., 2013. Flower diversity and angiosperm diversification. In J. L. Riechmann & F. Wellmer, eds. Flower Development: Methods and Protocols. New York: Springer New York, pp. 85–102.

Stark, P., 1918. Die Blütenvariationen der Einbeere. Zeitschrift für induktive Abstammungsund Vererbungslehre 19: 241–303.

Stebbins, G.L., Jr., 1974. Flowering Plants: Evolution Above the Species Level, Cambridge: Harvard University Press.

Stevens, P.T., C.A. Huether, and T.K. Wilson. 1972. Apical size in the determination of corolla lobe number in Linanthus androsaecus ssp androsaecus. American Journal of Botany 59: 989–992.

Strauss, S.Y., and J.B. Whittall. 2006. Non-pollinator agents of selection on floral traits. Ecology and evolution of flowers.

Tooke, F., and N.H. Battey. 2000. A leaf-derived signal is a quantitative determinant of floral form in Impatiens. The Plant Cell 12: 1837–1847.

Van Natto, A.C. and C.G. Eckert. Genetic and conservation significance of populations at the polar vs. equatorial range limits of the Pacific coastal dune endemic Abronia umbellata (Nyctaginaceae). Conservation Genetics 23: 255–269.

Vlot, E.C., W.H.J. van Houten, S. Mauthe, and K. Bachmann. 1992. Genetic and Nongenetic Factors Influencing Deviations from Five Pappus Parts in a Hybrid between Microseris douglasii and M. bigelovii (Asteraceae, Lactuceae). International Journal of Plant Sciences.

Warren, J. 2009. Extra petals in the buttercup (Ranunculus repens) provide a quick method to estimate the age of meadows. Annals of Botany 104: 785–788.

